# Bayesian Tests for Random Mating in Polyploids

**DOI:** 10.1101/2022.08.11.503635

**Authors:** David Gerard

## Abstract

Hardy-Weinberg proportions (HWP) are often explored to evaluate the assumption of random mating. However, in autopolyploids, organisms with more than two sets of homologous chromosomes, HWP and random mating are different hypotheses that require different statistical testing approaches. Currently, the only available methods to test for random mating in autopolyploids (i) heavily rely on asymptotic approximations and (ii) assume genotypes are known, ignoring genotype uncertainty. Furthermore, these approaches are all frequentist, and so do not carry the benefits of Bayesian analysis, including ease of interpretability, incorporation of prior information, and consistency under the null. Here, we present Bayesian approaches to test for random mating, bringing the benefits of Bayesian analysis to this problem. Our Bayesian methods also (i) do not rely on asymptotic approximations, being appropriate for small sample sizes, and (ii) optionally account for genotype uncertainty via genotype likelihoods. We validate our methods in simulations, and demonstrate on two real datasets how testing for random mating is more useful for detecting genotyping errors than testing for HWP (in a natural population) and testing for Mendelian segregation (in an experimental S1 population). Our methods are implemented in Version 2.0.2 of the hwep R package on the Comprehensive R Archive Network https://cran.r-project.org/package=hwep.

## 1 Introduction

One reason researchers often test for Hardy-Weinberg proportions (HWP) is to evaluate the presence of random mating [Waples, 2015]. At autosomal loci in diploids, one or two generations of random mating, for monoecious and dioecious species, respectively, are sufficient to reach HWP [Gillespie, 2004]. Thus, for diploids, random mating and HWP are effectively equivalent hypotheses, and so use the same statistical procedures. However, in autopolyploids, organisms with more than two sets of homologous chromosomes, HWP are only approached asymptotically [Haldane, 1930], and so random mating and HWP are separate hypotheses. To put notation to these hypotheses, let ***q*** = (*q*_0_, *q*_1_, …, *q*_*K*_) be the genotype frequencies of a *K*-ploid population at a biallelic locus with allele frequency 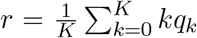. That is, *q*_*k*_ is the proportion of the population with *k* copies of the minor allele. Then the HWP hypothesis states that, in the absence of double reduction [Huang et al., 2019], the genotype frequencies follow binomial proportions,

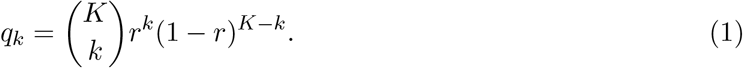

While, under the hypothesis of random mating, we have

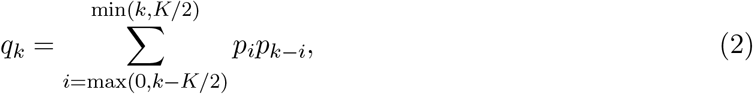

where ***p*** = (*p*_0_, *p*_1_, …, *p*_*K*/2_) are the *gamete* frequencies of the population. That is, *p*_*k*_ is the proportion of gametes in the population that have *k* copies of the minor allele. Equation (2) is called a “discrete linear convolution”, and can be represented concisely by ***q*** = ***p*** ∗ ***p***. Equation (2) arises because the genotype of an individual is the sum of the genotypes of two independent gametes (one from each parent), and the distribution of the sum of independent random variables is a convolution of the distributions of the summands. Hypotheses (1) and (2) are clearly distinct, which indicates the necessity to create separate testing procedures to evaluate them.

Testing for binomial proportions (1) is a fairly straightforward statistical problem, and several researchers have developed permutation approaches to do so effectively, particularly for small counts [Hardy and Vekemans, 2002, Meirmans and Van Tienderen, 2004]. By contrast, the only approaches that test for random mating (2) heavily rely on asymptotic approximations [Sun et al., 2020, Gerard, 2022b]. (Though, Matoka Nana [2023] developed exact tests for random mating in autopolyploids, but these tests were found to be far too conservative for practical use). Such asymptotic approximations are particularly fraught in autopolyploids where many genotypes are possible, and sample sizes are typically small. Thus, in most applications, there will be genotypes with few/no individuals, limiting the applicability of asymptotic approaches. The current approaches also do not account for genotype uncertainty, a pervasive issue in polyploid data [Gerard et al., 2018, Gerard and Ferrão, 2019], and one that limits the real-world applicability of current methods.

Furthermore, the only approaches that test hypothesis (1) use frequentist procedures. Sun et al. [2020] developed a test for random mating in tetraploids (*K* = 4) that finds the MLEs under model (2) via an EM algorithm, and uses these estimated genotype frequencies to run a chi-squared test. Sun et al. [2020] called this a “A Gamete-Based aHWE [asymptotic Hardy-Weinberg equilibrium] Test”, but this is incorrect as it is a test for random mating, not HWP. Sun et al. [2020] also uses an incorrect degrees of freedom for their test [Gerard, 2022a]. Gerard [2022b] generalized this EM algorithm to every ploidy (*K* ≥ 4) and implemented likelihood ratio tests for random mating for every ploidy. One issue with frequentist procedures is that, for increasing sample size, the *p*-value threshold should decrease to provide consistency under the null [Wakefield, 2009], but this is rarely done.

Bayesian approaches provide solutions to all of the above issues. Small sample sizes are not an issue for Bayesian methods as all inference is conditional on the observed data, not on hypothetical realizations of data which most frequentist approaches approximate via asymptotics. Bayesian methods provide a natural foundation to integrate over genotype uncertainty. And, whereas frequentist *p*-values should have lower significance levels for larger sample sizes so that both type I and type II error rates decrease to zero, Bayes factors automatically calibrate themselves with increasing sample size [Shoemaker et al., 1998, Wakefield, 2009]. The appeal of the Bayesian approach can be evidenced by the large number of methods in the diploid literature that implement Bayesian tests for HWP [Altham, 1971, Pereira and Rogatko, 1984, Lindley, 1988, Shoemaker et al., 1998, Montoya-Delgado et al., 2001, Consonni et al., 2008, Wakefield, 2010, Consonni et al., 2011, Puig et al., 2017, 2019].

The goal of this paper is to produce and implement Bayesian tests for hypotheses

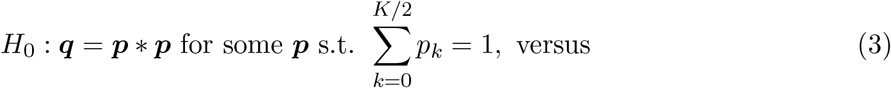

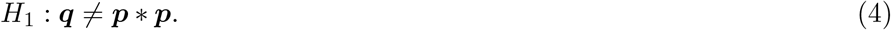

When genotypes are known, we implement a Bayesian test via an efficient Gibbs sampler that uses a novel data augmentation scheme (Section 2.1). However, genotypes are rarely known in polyploid organisms, and so we account for genotype uncertainty using genotype likelihoods (Section 2.2). Using simulations, we explore the behavior of our tests when genotypes are known (Section 3.1), before exploring the performance of our procedures under genotype uncertainty (Section 3.2). We demonstrate the utility of our methods on both a natural population of tetraploid white sturgeon (Section 3.3) and an experimental population of hexaploid sweet potatoes (Section 3.4), where we show that competing methods are less useful, particularly at detecting genotyping errors.

## 2 Materials and methods

### 2.1 Bayesian test with known genotypes

In this section, we will create a Bayesian test for random mating when genotypes are known. We will limit ourselves to summaries in the main text, leaving more technical details to Appendix S1. Let *x*_*k*_ be the number of individuals with genotype *k* ∈ {0, 1, …, *K*}, which we collect into the vector ***x*** = (*x*_0_, *x*_1_, …, *x*_*K*_). Then, given genotype frequencies ***q***, we have that

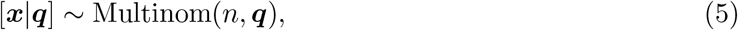

where Multinom(*n*, ***q***) denotes the multinomial distribution with size 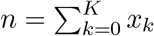 and probability vector ***q***. Our goal is to use likelihood (5) to test hypotheses (3)–(4). We will do so by calculating a Bayes factor [Kass and Raftery, 1995], the ratio of the marginal likelihoods under the null and alternative,

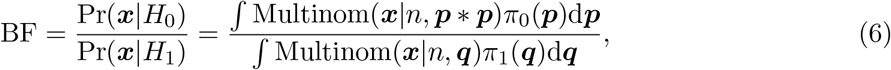

where *π*_0_(·) and *π*_1_(·) denote prior densities under the null and alternative, respectively, and Multinom(***x***|*n*, ***q***) is the multinomial likelihood of genotype counts ***x***, size *n*, and probability vector ***q***. In this manuscript, we will assume that these prior densities are both Dirichlet (Section 2.3) with concentration hyperparameters ***α*** under the null and ***β*** under the alternative,

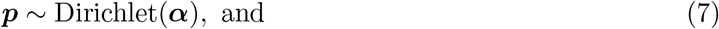

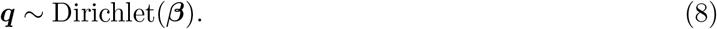

Smaller Bayes factors provide more evidence for *H*_1_, and larger Bayes factors provide more evidence for *H*_0_.

We will need to separately calculate the integrals in the denominator and numerator of (6). The denominator in (6) is a standard calculation. The integral,

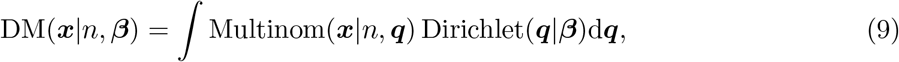

results in a Dirichlet-Multinomial distribution [Mosimann, 1962]. This has a known form (S1) and is easily computable.

To calculate the marginal likelihood in the numerator of (6), we will need a data augmentation scheme to represent the distribution of ***x*** ∼ Multinom(*n*, ***p***∗***p***), as this will allow us to use conjugacy to derive a closed form for Pr(***x***|*H*_0_). Let us begin by considering how random mating arises via the union of two randomly sampled, unobserved gametes. Let ***y*** = (*y*_0_, *y*_1_, …, *y*_*K*/2_) be the gamete counts, where *y*_*j*_ is the number of gametes in our sample with genotype *j*. Random mating implies that these gametes are selected with replacement according to the gamete frequencies, i.e. ***y*** ∼ Multinom(2*n*, ***p***). These gametes then randomly pair to form the next generation of offspring, and we denote the counts of pairings by the matrix ***A*** = (*a*_*ij*_), where *a*_*ij*_ is the number of individuals who were formed from gametes with dosages *i* and *j* with *i* ≤ *j*. The number of offspring with genotype *k*, which we denote by *x*_*k*_, is derived by summing over the cells of ***A*** such that *k* = *i* + *j*,

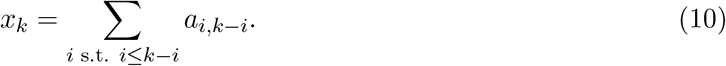

It is easy to show that ***y*** is also a function of ***A***,

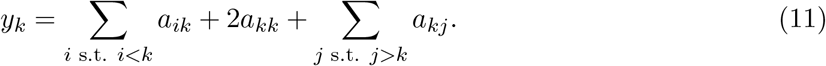

See Figure 1 for a visualization of the connections between ***A, x***, and ***y***.

**Figure 1:**
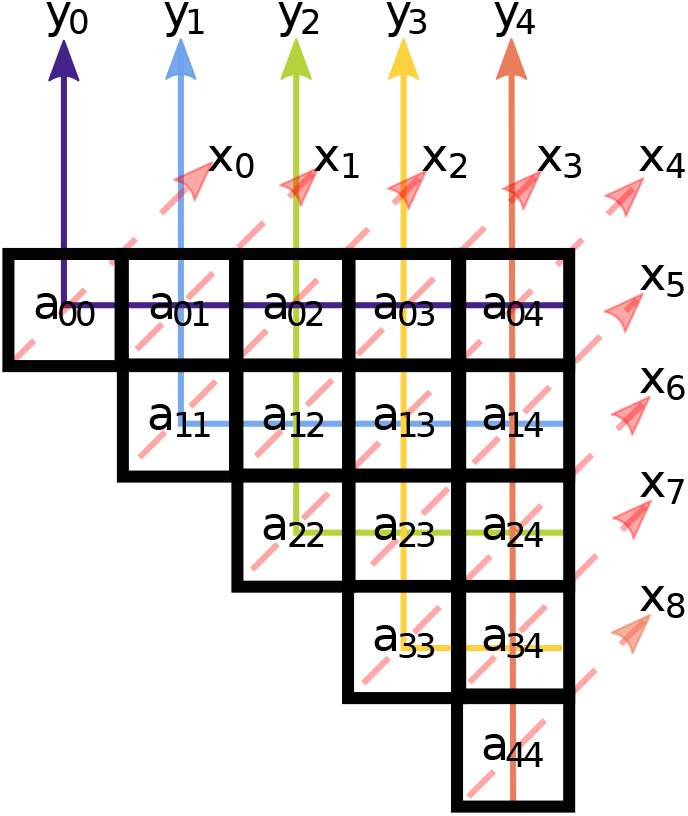
Visualization of data augmentation from (10) and (11) for an octoploid species (*K* = 8). The elements of ***A*** are in each cell. The *x*_*k*_’s are the sums of the anti-diagonal elements, indicated by the dashed red lines. The *y*_*k*_’s are the sums of the color-coded solid lines.

We will write the probability mass function (PMF) of ***x*** by marginalizing over the unobserved quantities ***A*** and ***y***. We know the PMF of [***y***|***p***] is Multinom(***y***|2*n*, ***p***). Further, the PMF of [***A***|***y***] is a known form (S6) [Levene, 1949], which we will denote by *f*(***A***|***y***). Also, since ***y*** is a function of ***A*** (11), which we denote by ***y***(***A***), the PMF of ***A*** may be written as *f*(***A***|***y***(***A***)) Multinom(***y***(***A***)|2*n*, ***p***). We may use this to marginalize over ***A*** to derive the following identity,

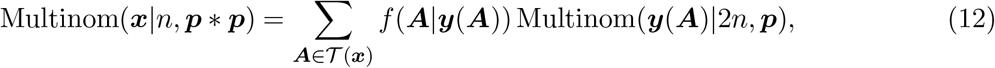

where 𝒯(***x***) is the set of ***A***’s that are consistent with ***x*** (Figure 1).

Identity (12) allows us to derive the marginal likelihood under the null in closed form when using a Dirichlet prior over ***p***. Namely.

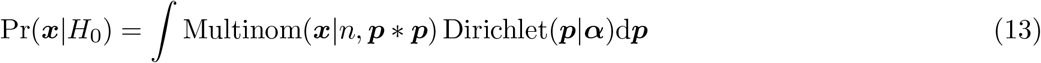

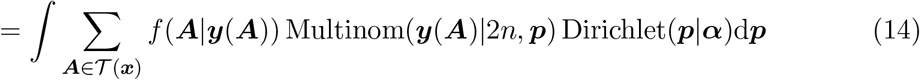

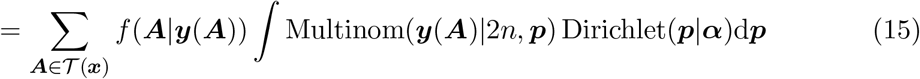

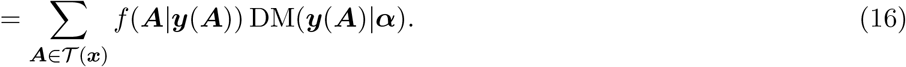

We describe in Appendix S1 how to sum over (16).

Summation (16) is efficient to calculate for tetraploids, and is possible to calculate for small-to-moderate sample sizes of hexaploids. However, for larger ploidies the summation (16) becomes too computationally intensive for all but the smallest of sample sizes. Therefore, in Appendix S2, we derive a Gibbs sampler [Gelfand and Smith, 1990] to sample from the posterior distribution of [***p***|***x***] using the same data augmentation scheme as in Figure 1. In this Gibbs sampler, we iteratively sample [***A***|***p, x***] and [***p***|***A***], and the resulting marginal samples of ***p*** are from the distribution of [***p***|***x***]. The full Gibbs sampler is summarized in Procedure S1. After obtaining the posterior samples from [***p***|***x***], we can estimate the marginal likelihood by the method of Chib [1995].

### 2.2 Bayesian test using genotype likelihoods

Due to the higher number of possible dosages, polyploids are more susceptible than diploids to data variability and data biases, and thus exhibit higher amounts of genotype uncertainty [Gerard et al., 2018, Gerard and Ferrão, 2019, Gerard, 2021a,b]. A common way to account for this is through the use of genotype likelihoods [Li, 2011]. A nice feature of Bayesian methodology is its ability to easily adapt to additional variability by integrating over it. Here, we develop Bayesian testing procedures for random mating that account for genotype uncertainty using genotype likelihoods.

It is possible to modify our Gibbs sampler from Section 2.1 to account for genotype likelihoods via a data augmentation scheme where we introduce latent genotypes that we sample over (Appendix S3). This approach works well for small ploidies and small sample sizes. However, we have found that this approach produces poor mixing (i.e., it inefficiently explores the posterior) when both the ploidy *K* and the sample size *n* is large (Section 3.2), and thus requires a large number of samples to produce accurate results. We have therefore opted to use the Stan programming language [Stan Development Team, 2022a,b], which uses Hamiltonian Monte Carlo, to more efficiently explore the posterior distribution in fewer samples by using gradient information [see Betancourt, 2018, for an accessible introduction]. Marginal likelihoods and Bayes factors may then be calculated via bridge sampling [Meng and Wong, 1996, Gronau et al., 2017, 2020].

Let us discuss the model implementation. Let *ℓ*_*ik*_ be the genotype likelihood for individual *i* = 1, 2, …, *n* for genotype *k* = 0, 1, …, *K*. That is, *ℓ*_*ik*_ is the probability of the data (sequencing, microarray, or otherwise) for individual *i* given that the genotype for that individual is *k*. Genotype likelihoods are provided by many genotyping programs [McKenna et al., 2010, Voorrips et al., 2011, Serang et al., 2012, Gerard et al., 2018, Clark et al., 2019, Gerard and Ferrão, 2019]. Then, given these genotype likelihoods, we have the likelihood for these data is

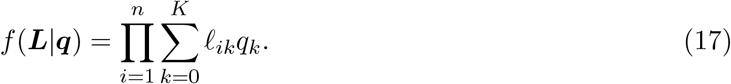

Under the alternative, the prior over ***q*** is (8), while under the null, the prior over ***q*** = ***p***∗***p*** is induced from (7). We run two Markov chains, one using prior (8) and one using prior (7), estimating the marginal likelihoods of each via bridge sampling, and from there calculating the Bayes factor. These implementations in Stan are relatively straightforward so we omit the specific details. The Stan files may be found in the source code for our hwep R package (https://cran.r-project.org/package=hwep).

### 2.3 Prior elicitation

Here, we describe our default priors. For convenience, as in Sections 2.1 and 2.2, we assume that the gamete frequencies are *a priori* Dirichlet(***α***) under the null, and the genotype frequencies are *a priori* Dirichlet(***β***) under the alternative, and our focus will be on selecting appropriate values of ***α*** and ***β***.

To motivate our choice for ***α***, we note that at HWP, in the absence of double reduction, we have for autopolyploids that the *p*_*i*_’s are binomial proportions for some allele frequency *r* [Haldane, 1930]. That is,

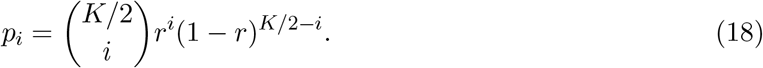

For natural populations, it is reasonable to place a uniform prior over *r*. Matching moments with the Dirichlet, we set *α*_*i*_ to be proportional to the expected value of (18) when *r* is uniformly distributed. We have that

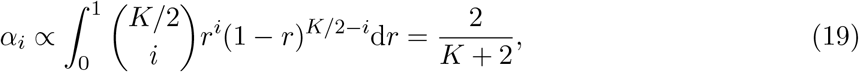

where the calculation in (19) uses the kernel of the Beta density. Thus, each *α*_*k*_ should have the same value *a priori*, and this motivates us to take

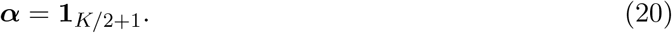

That is, ***p*** is uniformly distributed over the *K*/2 unit simplex. This might appear to be just the naive prior one would choose as “uninformative”, but we have also motivated it by placing an uninformative prior on the allele frequency of a randomly mating natural population at HWP. Though we have motivated it using HWP, it is important to note that our methods *do not* assume HWP.

We choose our prior for ***β*** conditional on our choice of ***α***. Our elicitation of ***β*** makes the following key assumption: *samples of size one are completely uninformative on the status of random mating*. This approach is conceptually similar to that of Smith and Spiegelhalter [1980] and Spiegelhalter and Smith [1982], except they chose their datasets that are most favorable to the null (since they worked in a continuous data paradigm) while we choose all possible datasets where *n* = 1. To specify this prior, let ***e***_*k*_ be a 1-of-(*K* + 1) vector that is 1 at position *k* and is 0 elsewhere (so ***e***_*k*_ is the value of ***x*** when *n* = 1), then our assumption is that

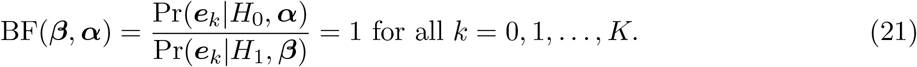

We can show that, given ***α*** = **1**_*K*/2+1_, for any choice of scaling factor *c >* 0, the following choice of ***β*** will result in property (21),

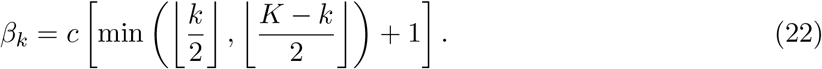

Equation (22) is merely a constant *c* multiplied by the number of elements in ***A*** that correspond to each *x*_*k*_ (Lemma S1). We prove this result in Corollary S1 in Appendix S4.

By default, we choose a value of *c* such that the total sum of ***β*** is the same as **1**_*K*+1_. This intuitively means that we place the same overall weight on the prior as a uniform distribution would. When ***α*** = **1**_*K*/2+1_, this results in default values of ***β***,

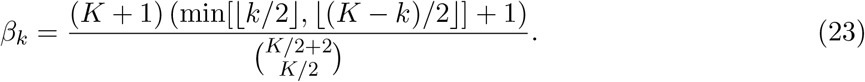

The default values of ***β*** for even ploidies less than or equal to 12 are presented in Table S1.

It is important to note that priors can impact Bayes factors for even large sample sizes [Kass and Raftery, 1995], and so choosing reasonable default priors is extremely important. However, because our priors are proper, have support over the entire parameter space, and the models are nested, the Bayes factors are guaranteed to be consistent for large sample sizes [O’Hagan, 1994, Section 7.52]. That is, the Bayes factor goes to infinity if the null is true and goes to zero if the alternative is true. Thus, for large sample sizes we will at least make the correct statistical decision, even if the Bayes factors do not accurately describe the odds of the null to the alternative that are given by the data. We evaluate prior sensitivity in Appendix S5.

## 3 Results

### 3.1 Performance when genotypes are known

We can verify that the Bayes factors increase under the null, and decrease under the alternative, as the sample size increases. To do this, we simulated genotype counts for ploidies *K* ∈ {4, 6, 8} with *n* ∈ {10, 100, 1000} individuals under two scenarios. The first scenario was where the null was satisfied with ***p*** = **1**_*K*/2+1_/(*K*/2 + 1) and ***q*** = ***p*** ∗ ***p***, and the second scenario was where the alternative was satisfied with ***q*** = **1**_*K*+1_/(*K* + 1). For each scenario, we generated genotype counts via ***x*** ∼ Multinom(*n*, ***q***). Each replication, we obtained the Bayes factor from Section 2.1 and the *p*-value from Appendix C of Gerard [2022b] (a likelihood ratio test for random mating). For each combination of parameters, we replicated these simulations 1000 times.

The results are in Figure 2. There, we see that the Bayes factor is consistent under both the alternative and null scenarios. We also plot the computation time in Figure S1 under these scenarios. We see that, when using simulation, the Bayesian approach has about the same computation time for any ploidy and sample size. The Bayesian test (which is *O*(1)) is even slightly faster than the likelihood ratio test from Appendix C of Gerard [2022b], but both are still fast enough for genome-wide applications. Perhaps the biggest disadvantage of the likelihood ratio test is that the *p*-values, under the null, are not uniformly distributed for small sample sizes (Figure 3) because of its reliance on asymptotics. The Bayesian approach does not depend on asymptotics and so may be used at any sample size.

**Figure 2:**
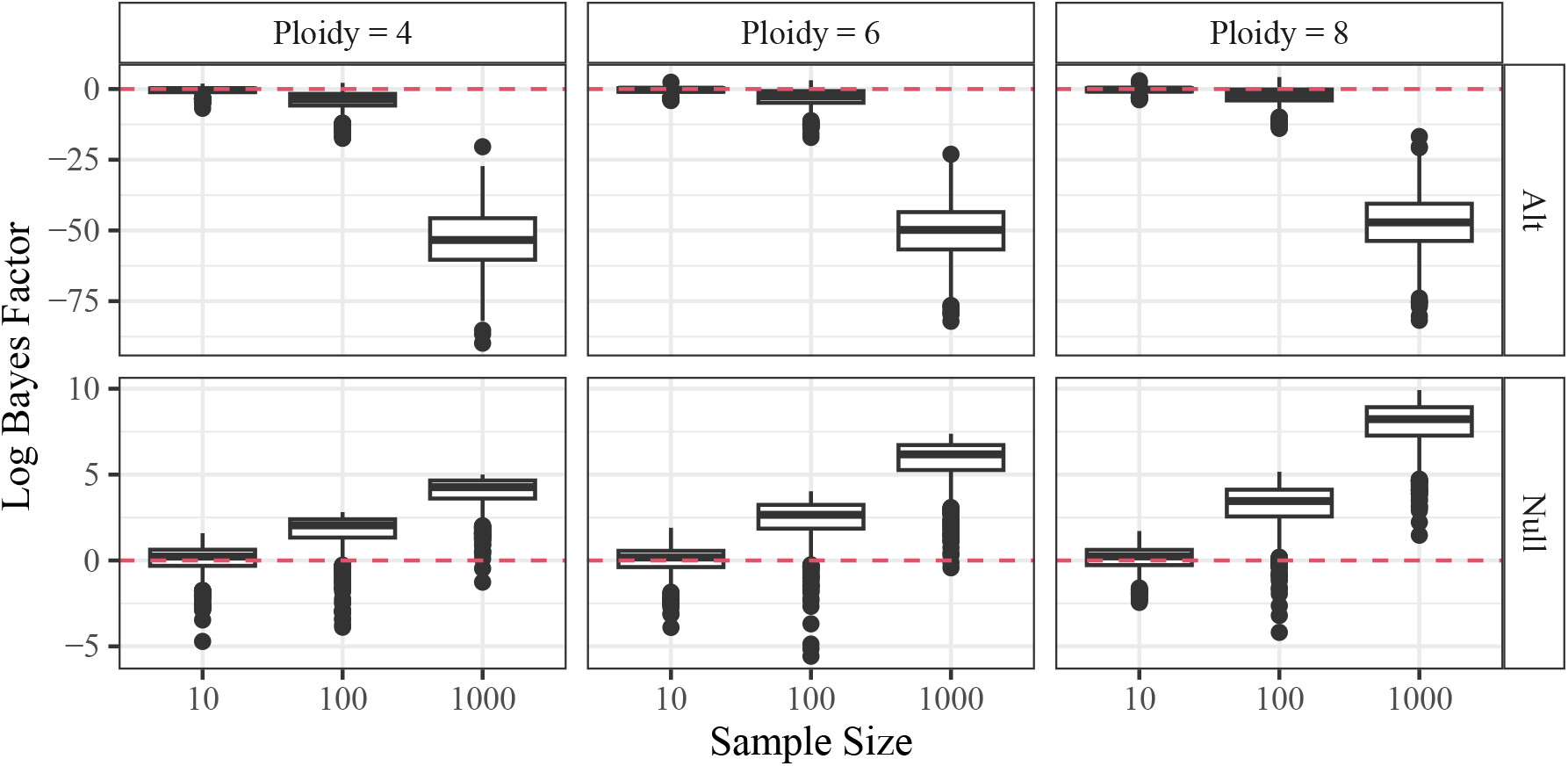
Log Bayes factors (*y*-axis) at different sample sizes (*x*-axis) at different ploidies (column facets) when either the null is true (“Null”) or the alternative is true (“Alt”). The Bayes factor increases gradually with sample size when the null is true and decreases rapidly with sample size when the alternative is true.

**Figure 3:**
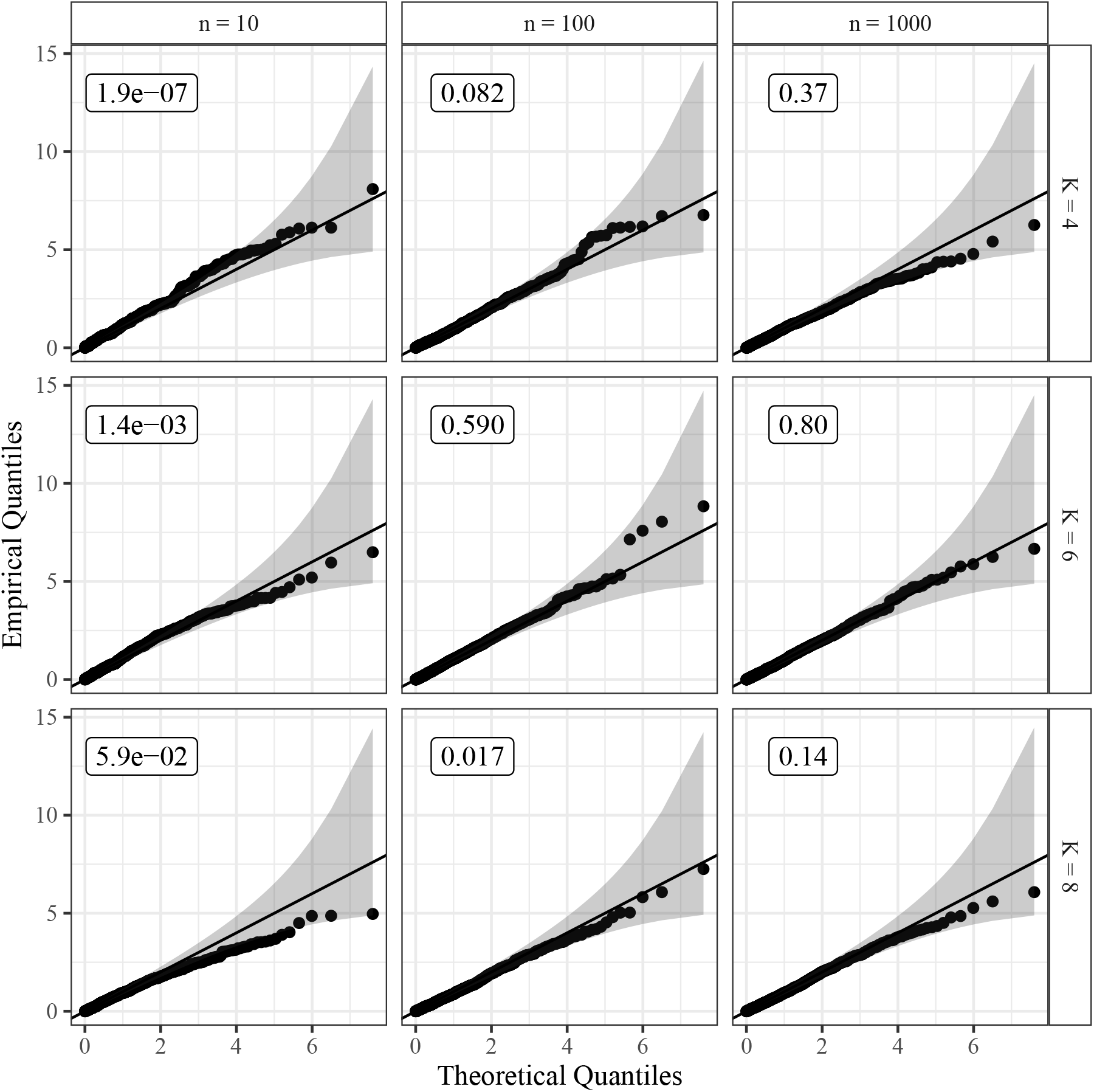
QQ-plots of *p*-values of the likelihood ratio test for random mating against the uniform distribution, on the *−* log scale, with the 95% simultaneous confidence bands from Aldor-Noiman et al. [2013], stratified by ploidy (row facets) and sample size (column facets). These *p*-values come from the simulations of Section 3.1 when the null is satisfied, and so should follow a uniform distribution. The numbers in the top-left of each plot are the *p*-values from a Kolmogorov-Smirnov test against the null of uniformity. Only for larger sample sizes are the *p*-values appropriately calibrated.

In Appendix S6, we consider the behavior of our tests when the sample size is very small. We show that the range of possible Bayes factors is limited when samples are ∼ 5 (Figures S2 and S3, and Table S2). At such sample sizes, only moderate evidence is possible either for or against the random mating hypothesis. Of course, this is a property of the data and this limitation also applies to frequentist approaches.

To evaluate the sensitivity of our methods to prior selection, we ran a prior sensitivity analysis in Appendix S5. There, we confirm that the Bayes factors are consistent for various prior selections, and we demonstrate that the Bayes factors are relatively stable to moderate deviations from our default priors (Figure S4).

### 3.2 Performance under genotype uncertainty

To explore the performance of our approaches under genotype uncertainty, we incorporated genotype likelihoods into the simulation study from Section 3.1. For each replication, using the same simulation settings as in Section 3.1, we simulated read counts using rflexdog() from the `updog R` package [Gerard et al., 2018, Gerard and Ferrão, 2019] at a read depth of 10, a sequencing error rate of 0.01, an overdispersion parameter of 0.01, and no allelic bias. From these read counts, we then used `updog` (with the general categorical prior class) to generate genotype likelihoods and genotype posteriors. We used these genotype likelihoods to run the Bayesian test from Section 2.2. To compare this approach to that of Section 2.1, we need an estimate of ***x***. We did this by two methods: (i) summing up the posterior probabilities for each genotype across individuals and then rounding these sums to the nearest integers, and (ii) obtaining the posterior mode genotype for each individual and tabulating the genotype counts.

The results are presented in Figure S5. We see there that only the genotype likelihood approach from Section 2.2 is consistent, with the Bayes factor increasing with the sample size when the null is satisfied. Assuming known genotypes, the method of Section 2.1 appears to be anti-conservative in the presence of genotype uncertainty, generally providing Bayes factors that are too small, when using either approach to generate ***x***. This indicates the need to account for genotype uncertainty in this testing situation to provide accurately calibrated test statistics. The likelihood ratio test for random mating from Appendix C of Gerard [2022b], which assumes known genotypes, also behaves very poorly in the presence of genotype uncertainty (Figure S6).

To evaluate the sensitivity of our methods to the calibration of the genotype likelihoods, we altered the conditions of `updog` when genotyping the simulated read-counts. We either estimated the likelihood parameters (sequencing error rate, overdispersion, and allelic bias) or fixed them at their known values. When estimating the likelihood parameters, the prior class can greatly affect the estimates [Gerard and Ferrão, 2019] (the prior class has no effect on the genotype likelihoods when the likelihood parameters are known), and so when estimating these parameters we either used the general categorical prior class, the proportional normal prior class [Gerard and Ferrão, 2019], or the discrete uniform prior. The ideal scenario is using known likelihood parameters. The typical scenarios are using either the proportional normal prior or the general categorical prior and estimating the likelihood parameters. The most poorly calibrated scenario is using the discrete uniform prior while estimating the likelihood parameters [Gerard et al., 2018]. The results are in Figure S7. There, we see that all approaches to generating the genotype likelihoods yield consistent results, indicating that our methods are very robust to genotype likelihood miscalibration. Though, using the uniform prior or the general categorical prior class does perform worse.

We also evaluated the Gibbs sampler from Appendix S3 in these simulations. The number of iterations in the Gibbs sampler was chosen to have the same run-time as the Stan approach when *K* = 8 and *n* = 1000 (Figure S8). The Bayes factors calculated using the Gibbs sampler and those calculated using Stan are mostly the same except when both the sample size and ploidy are large (Figures S9–S10). The differences are due to poor mixing in the Gibbs sampler, as this approach is not consistent for *K* = 8 (Figure S11), at least given the number of sampling iterations used. This demonstrates the benefits of using Stan, at least for large sample sizes and large ploidies, to improve mixing.

### 3.3 A natural population

In this section, we analyze the tetraploid white sturgeon (*Acipenser transmontanus*) data from Delomas et al. [2021]. These data were collected from a central California caviar farm with broodstock from the Sacramento river, and so likely exhibit random mating. We genotyped the *n* = 19 individuals at 325 loci using `updog` [Gerard et al., 2018, Gerard and Ferrão, 2019]. These data were sequenced to a very high average read depth of 5342, and so it is reasonable to take genotypes at most loci as known. We ran our Bayesian test (Section 2.1) and the likelihood ratio test for random mating from Gerard [2022b] on these loci.

Figure 4 contains a histogram of the log Bayes factors. We see there that the mode of the log Bayes factors is above 0, with most log Bayes factors near or slightly above zero. This provides strong evidence that the population is undergoing random mating. There are just three loci that appear to deviate from this trend (Table S3).

**Figure 4:**
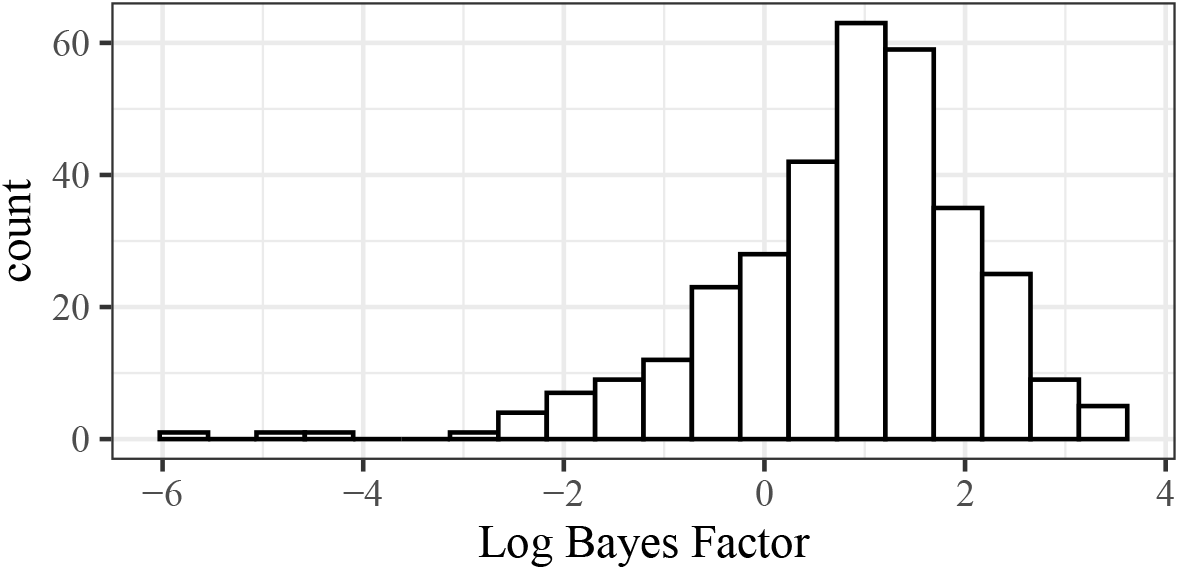
Histogram of log Bayes factors applied to all 325 loci from the white sturgeon data from Section 3.3. We see that most log Bayes factors are near or slightly above zero, indicating strong evidence that the population is undergoing random mating.

The test for random mating immediately reveals its use as, looking deeper at the three loci with the smallest Bayes factors, it appears that all three are a result of genotyping errors. Let us take, as an example, the SNP with the smallest Bayes factor, Atr_20529-52. When we fit `updog` on these data, there are two modes of the log-likelihood (Figure 5). The maximum likelihood estimate results in ***x*** = (0, 6, 2, 1, 10) (log-likelihood of -90) while a smaller local mode results in ***x*** = (0, 0, 6, 3, 10) (log-likelihood of -105). The local mode was found by starting the algorithm of `updog` at a different value of the bias parameter. This local mode estimates the allelic bias (the ratio of mapping probabilities of the minor and major alleles) to be 2.2, which is large but not too out of the ordinary for these data (Figure S12). Interestingly, when we run our Bayesian test for random mating on this local mode with ***x*** = (0, 0, 6, 3, 10), we get a log Bayes factor of 1.1, which is in alignment with the rest of these data (Figure 4). Thus, it is likely that this SNP was a genotyping error. When we do the same analysis with the SNPs with the next two smallest Bayes factors, Atr_33123-54 and Atr_26310-63, we also see that the global modes are likely genotyping errors and the local modes likely provide the correct genotype estimates (Figures S13–S14). This demonstrates the practical usefulness of our approach.

**Figure 5:**
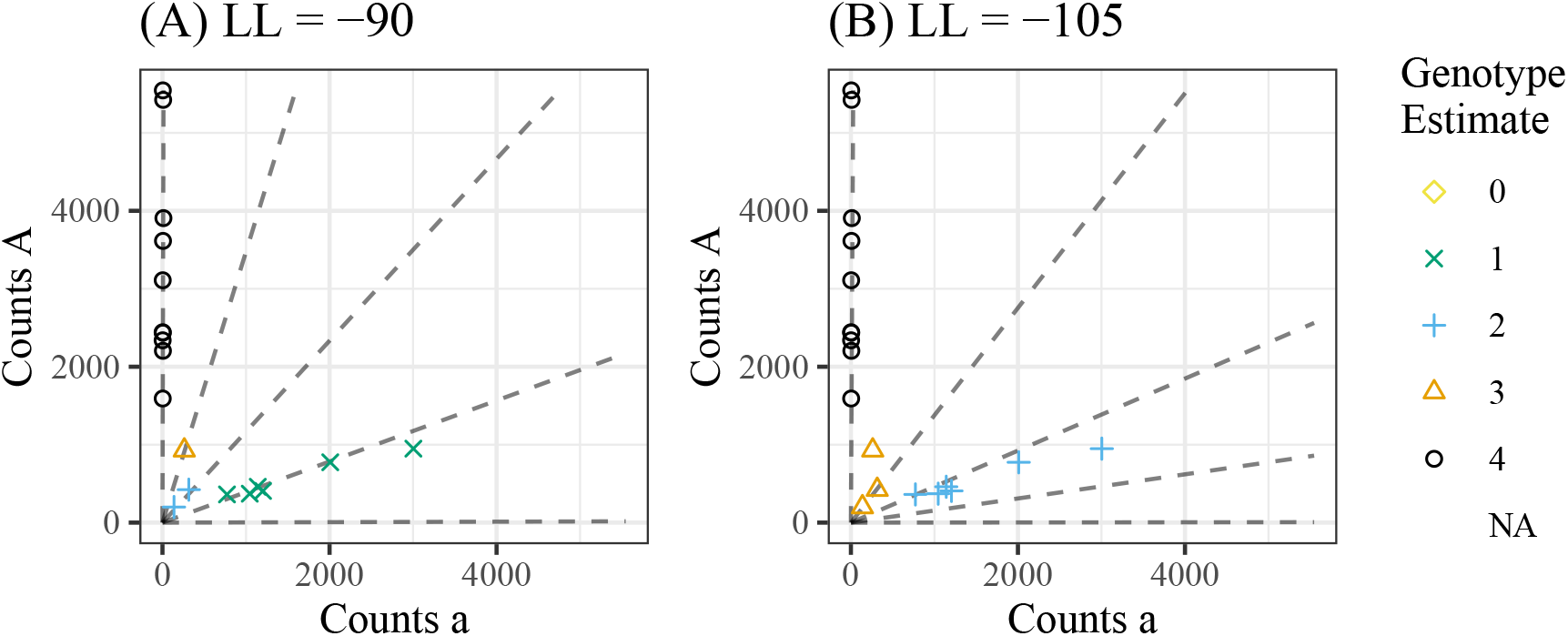
Genotype plots [Gerard et al., 2018] for two fits on SNP Atr_20529-52 from the white sturgeon data of Section 3.3. Each point is an individual, the color and shape of which indicate the estimated genotype under the two fits. We have the number of alternative reads on the *x*-axis and the number of reference reads on the *y*-axis. The lines radiating from the origin indicate the expected number of reference/alternative reads of a given genotype under that given fit. The left plot **(A)** is the global mode with a log-likelihood of -90, and the right plot **(B)** is a local mode with a log-likelihood of -105. The local mode exhibits high allelic bias, which can be seen by the lines clustering towards the *x*-axis. The test for random mating using the global mode of ***x*** = (0, 6, 2, 1, 10) yields a log Bayes factor of -5.9, while the test for random mating using the local mode of ***x*** = (0, 0, 6, 3, 10) yields a log Bayes factor of 1.1. The global mode is thus likely a genotyping error.

The likelihood ratio test for random mating from Gerard [2022b] produces log *p*-values that are linearly associated with the log Bayes factors, particularly in the tail (Figure S15). However, standard frequentist approaches would indicate that no loci exhibit strong deviations from random mating. For example, the smallest three *q*-values [Storey, 2003] are 0.06, 0.17, and 0.17, which many researchers would not raise an eyebrow to since they are all above 0.05.

Tests for HWP are less useful for these data. The *p*-values, using the *U*-statistic based test from Gerard [2022b], are highly non-uniform (Figure S16). Indeed, using these *p*-values, the `qvalue` package [Storey, 2003] estimates that 53% of loci do not satisfy the HWP hypothesis, with 14 loci being rejected at the 0.05 FDR level. This indicates that, though these data exhibit random mating, there are some severe deviations from HWP, likely caused by unobserved relatedness between individuals. Indeed, when we calculate the pairwise relatedness measure of Ashraf et al. [2016] (the autopolyploid extension of VanRaden [2008]), using the `AGHmatrix R` package [Amadeu et al., 2020], we see two distinct clusters of closely related individuals (Figure S17). This limits the usefulness of HWP tests for detecting genotyping errors.

### 3.4 An S1 population

Here, we analyze the S1 population of 142 autohexaploid sweet potatoes (*Ipomoea batatas*) from Shirasawa et al. [2017]. An S1 population consists of a single generation of selfing from the same parent. It might seem strange, but random mating is fulfilled in S1 populations because (i) all offspring gametes are drawn independently and identically distributed from the distribution of gametes produced by the parent, and (ii) these gametes are randomly paired during fertilization. Thus, random mating should be fulfilled at most loci in this dataset, though some effects (e.g., inbreeding depression) might cause localized deviations from random mating.

Data were prepared and filtered as in Gerard [2022b]. That is, SNPs were selected that had allele frequencies between 0.1 and 0.9, average read depths of at least 100, and no more than 50% missing data. This resulted in 5074 SNPs. Genotype likelihoods were obtained by `updog` [Gerard et al., 2018, Gerard and Ferrão, 2019]. We used these genotype likelihoods to fit the Bayesian test from Section 2.2. The numbers of individuals with each dosage were estimated by summing posterior genotype probabilities across individuals and then rounding these sums. We used these estimated counts to fit the Bayesian test from Section 2.1 on these loci.

The Bayes factors calculated using either genotype likelihoods (Section 2.2) or assuming known genotypes (Section 2.1) are only moderately correlated (Figure S18), indicating that there is enough genotype uncertainty that we should only use the genotype likelihood approach for these data (Section 3.2). The log Bayes factors for these loci are almost all positive (Figure S19) indicating strong evidence of random mating, as it should for these data since random mating is indeed fulfilled.

We explored the 5 SNPs with log Bayes factors less than -5. Genotype plots for the `updog` fits of these SNPs are provided in Figure S20. `Updog` has its own measure of genotyping performance, the estimated proportion of individuals mis-genotyped, and yet all but one of these SNPs have reasonable values of this measure (Table S4), so would likely pass filters using it. The numbers of individuals with each genotype deviate strongly from that expected under Mendelian segregation (Table S4), indicating that these are, indeed, problematic SNPs.

Since this is an S1 population, it is possible to use this knowledge to specifically test for genotype frequencies that would result from Mendelian segregation. We could then compare this approach with our more general test for random mating. We created a Bayesian test for segregation distortion in S1 populations in Appendix S7 and applied this test to the sweet potato data. There was significant agreement on which SNPs satisfy the random mating assumption (Table S5). For SNPs that support the random mating hypothesis, the S1 test tends to produce larger Bayes factors, indicating greater discriminative ability, as expected (Figure S21). However, the S1 test provides many more SNPs that support the alternative hypothesis. Genotype plots of a random sample of these SNPs, where the S1 test indicates violations in the null and the random mating test indicates no violations, are presented in Figure S22. These SNPs appear to be relatively well behaved, except for two or three individuals in each SNP, where Mendelian segregation would predict those genotypes to be impossible. Indeed, those problematic individuals are the reasons why the S1 test is so keen on the alternative hypothesis, and if we remove them then the S1 test produces positive log Bayes factors (Table S6). If the goal of a test is to find SNPs where *any* individual might be mis-genotyped, then the S1 test might be appropriate. But it is very sensitive to minor violations in the genotype frequencies predicted by an S1 population, and so might be too finicky for applied work. By contrast, the random mating test only detects broad genotyping issues. We note, though, that these problematic individuals could possibly be a result of double reduction [Stift et al., 2010], in which case a more sophisticated method to test for S1 genotype frequencies in the presence of double reduction might produce better results.

## 4 Discussion

In this manuscript, we developed Bayesian tests for random mating in polyploids. Our methods are applicable both when genotypes are known and when there is high genotype uncertainty (e.g., due to low sequencing depth) using genotype likelihoods. The hypothesis of random mating is less strict than that of HWP in autopolyploids. Therefore one should prefer tests for random mating over those for autopolyploid HWP in many scenarios. We proposed using tests for random mating to detect genotyping errors, as is common practice with HWP tests in diploids [Hosking et al., 2004, Anderson et al., 2010, Winkler et al., 2014]. We demonstrated this application on two real datasets, in which the SNPs broadly follow random mating but not HWP, by detecting likely genotyping errors that were missed by other methods (including current frequentist approaches). Our Bayesian approaches are applicable for both small sample sizes (as they do not depend on asymptotics) and large sample sizes (as they are guaranteed to be consistent decision procedures).

Though this paper has focused on autopolyploids, our tests for random mating are also valid for allopolyploids. When allopolyploids exhibit random mating, they still produce genotype frequencies of the form ***q*** = ***p*** ∗ ***p***, the null hypothesis for our tests for random mating. However, whereas in Section 2.3 we elicited our prior for ***p*** by considering a natural population of autopolyploids at HWP, for allopolyploids we should update this prior by considering a natural population of allopolyploids at HWP. We derive our default priors when analyzing allopolyploid genomes in Appendix S4.1. We also describe in Appendix S8 how allopolyploid HWP can differ from random mating if there is dependence between the subgenomes, indicating that tests for random mating might be more appropriate for allopolyploids for many applications as well.

In addition to genotyping errors, random mating can be violated through various mechanisms, such as selection, inbreeding, and population structure. Both inbreeding and population structure can lead to widespread deviations from random mating across many SNPs, similar to the effects of inbreeding [Vieira et al., 2013] and population structure [Hao and Storey, 2019] on diploid populations, which cause broad deviations from HWP. On the other hand, selection, similar to genotyping errors, can result in localized deviations from random mating. Tests for random mating can be valuable in detecting any of these issues, just as tests for HWP are useful in detecting them in diploids [Waples, 2015].

Our methods assume that loci are biallelic. This is the most common scenario in practice, particularly in polyploid studies where reduced representation genotyping is often used [Baird et al., 2008, Elshire et al., 2011]. Extending random mating tests to multiallelic loci would greatly increase the complexity of the problem, and so we leave it for future work. However, we describe in Appendix S9 how to mathematically represent the hypothesis of random mating at multiallelic loci. The hypothesis of random mating mating (S74) contains many more parameters than (3), so one would have to be much more cautious about prior sensitivity, as any developed Bayes factor might not be as robust as our methods (Appendix S5).

In Section 2.3 we motivated our prior over ***p*** by considering a natural population at equilibrium, in the absence of double reduction. There, we assumed that such a natural population would have a uniform allele frequency distribution. The distribution of allele frequencies is called the “site frequency spectrum” and there has been significant work on deriving its properties [Fu, 1995, Griffiths and Tavaré, 1998, Wooding and Rogers, 2002, Polanski et al., 2003, Polanski and Kimmel, 2003, Birkner et al., 2013]. The distribution of the site frequency spectrum is a complicated function of population size and various aspects of data collection [Polanski and Kimmel, 2003]. If this information is known about a population, it might be possible to incorporate the theoretical expectations of the site frequency spectrum by modifying the expectation in (19), and thereby improve the prior elicitation. One would have to be careful about ascertainment bias caused, e.g., by using variants called from a previous sample, or the standard practice of filtering out SNPs with small minor allele frequencies. We do not explore these ideas further because our results in Appendix S5 indicate that any improvements from modifying the prior would be modest.

The random mating hypothesis could be used to improve genotyping methods. Many genotyping programs implement empirical Bayes approaches that estimate the genotype frequencies from a class of prior distributions before implementing a full Bayesian approach to genotyping [Voorrips et al., 2011, Serang et al., 2012, Blischak et al., 2018, Gerard et al., 2018, Clark et al., 2019, Gerard and Ferrão, 2019]. The choice of the class of prior distribution can significantly improve or harm the genotype calling performance [Gerard and Ferrão, 2019]. Our analysis in Section 3.3 suggests that we could improve genotyping performance by using (2) as the class of prior distributions, and doing so might have corrected the three genotyping errors we detected. Our methods here are intended to be agnostic to the genotyping method used, and so we leave implementing (2) as a prior class for genotyping as future work.

Most researchers are not used to the Bayesian approach to hypothesis testing, and so a common question is how to choose the threshold of the Bayes factor before one concludes violations in the random mating hypothesis? The canonical solution [Section 4.4.3 of Berger, 2013, e.g.] is for a researcher to elicit prior probabilities *π*_0_ and (1 *− π*_0_) for the null and alternative hypotheses, respectively, and losses *L*_*I*_ and *L*_*II*_ for type I and type II errors, respectively. One would then reject *H*_0_ if the Bayes factor is less than (1 *− π*_0_)*L*_*II*_/(*π*_0_*L*_*I*_). Eliciting these quantities can be difficult for most researchers. So our recommendations would be to (i) plot the Bayes factors and see if any loci are anomalous, (ii) in settings with few loci (∼ 100) use the thresholds from Kass and Raftery [1995] as a guide, and (iii) in GWAS settings with many loci use the threshold from Wakefield [2010] as a guide. For (ii) this would correspond to a log Bayes factor being less than -3 indicating strong evidence, and a log Bayes factor less than -5 indicating very strong evidence for violations of random mating. For (iii) this would correspond to a log Bayes factor less than -16 indicating very strong evidence for violations of random mating.

## Acknowledgements

This material is based upon work supported by the National Science Foundation under Grant No. 2132247.

Most analyses were performed using the R statistical language [R Core Team, 2022].

## Data accessibility

All methods discussed in this manuscript are implemented Version 2.0.2 of the hwep R package on the Comprehensive R Archive Network:

https://cran.r-project.org/package=hwep.

Scripts to reproduce all of the results of this manuscript are available on Zenodo:

https://doi.org/10.5281/zenodo.6993722.

All datasets used in this manuscript are open and available at:

https://www.doi.org/10.5061/dryad.crjdfn33r, and

http://sweetpotato-garden.kazusa.or.jp/

## Supplementary material

Additional details, figures, and tables are available in the Supplementary Material online.

## Author contributions

DG developed the methodology, wrote the software, implemented the study, and wrote the manuscript.

