## Supplementary Materials for "Bayesian Tests for Random Mating in Polyploids"

David Gerard

Department of Mathematics and Statistics, American University, Washington, DC, 20016, USA

**Abstract**

This document contains additional theoretical considerations, derivations, simulations, and figures to supplement the manuscript “Bayesian Tests for Random Mating in Polyploids”.

#### S1 Calculation of Bayes factors with known genotypes

In this section, we will describe how to obtain marginal likelihoods under both the null (3) and the alternative (4). The marginal likelihood under the alternative will be rather straightforward. For the null, we obtain a closed-form equation for the marginal likelihood. This is feasible to compute for tetraploids and small samples of hexaploids, but not for larger ploidies / sample sizes. Thus, we will then develop a Gibbs sampler to sample from the posterior distribution of gamete frequencies under the null, and from these samples we can estimate the marginal likelihood.

To calculate the Bayes factor (6), we will need the marginal likelihoods under both the null and the alternative. The marginal likelihood under the alternative is straightforward to calculate. Using prior (8) and likelihood (5), we have that $\boldsymbol{x}$ is Dirichlet-Multinomial [Mosimann, 1962], whose probability mass function (PMF) we will denote by

$$\Pr(\boldsymbol{x}|H_1) = \text{DM}(\boldsymbol{x}|n, \boldsymbol{\beta}) = \frac{\Gamma\left(\sum_{k=0}^{K} \beta_k\right)\Gamma(n+1)}{\Gamma\left(n + \sum_{k=0}^{K} \beta_k\right)} \prod_{k=0}^{K} \frac{\Gamma(x_k + \beta_k)}{\Gamma(\beta_k)\Gamma(x_k+1)}, \tag{S1}$$

where $\Gamma(\cdot)$ is the gamma function.

To calculate the marginal likelihood under the null, we will need the following data augmentation scheme to represent the multinomial likelihood with probability vector $\boldsymbol{p} * \boldsymbol{p}$. Each individual in our sample corresponds to two parental gametes, and we let $\boldsymbol{A} = (a_{ij})$, with $j \geq i$, denote the number of parental gamete pairs in our sample with genotypes $0 \leq i \leq j \leq K/2$. For example, if an individual's parents provided gametes with genotypes 2 and 0, then this would contribute one count to $a_{02}$. Let $\boldsymbol{y} = (y_0, y_1, \ldots, y_{K/2})$, where $y_k$ denotes the number of gametes in our sample with dosage $k$. Then we have the relationships

$$x_k = \sum_{i \text{ s.t. } i \leq k-i} a_{i,k-i}, \text{ and} \tag{S2}$$

$$y_k = \sum_{i \text{ s.t. } i<k} a_{ik} + 2a_{kk} + \sum_{j \text{ s.t. } j>k} a_{kj}. \tag{S3}$$

See Figure 1 for a visual description of these relationships. We can then represent the likelihood under the null in the following form

$$f(\boldsymbol{x}|n,\boldsymbol{p}) = \sum_{\boldsymbol{A}}\sum_{\boldsymbol{y}} f(\boldsymbol{x}|\boldsymbol{A})f(\boldsymbol{A}|\boldsymbol{y})f(\boldsymbol{y}|\boldsymbol{p}), \tag{S4}$$

where

$$f(\boldsymbol{x}|\boldsymbol{A}) = 1\left(x_k = \sum_{i \text{ s.t. } i\leq k-i} a_{i,k-i}\right), \tag{S5}$$

$$f(\boldsymbol{A}|\boldsymbol{y}) = \frac{n!\prod_{k=0}^{K/2} y_k!}{(2n)!\prod_{j\geq i} a_{ij}!}2^{\sum_{j>i} a_{ij}}, \text{ and} \tag{S6}$$

$$f(\boldsymbol{y}|\boldsymbol{p}) = \text{Multinom}(\boldsymbol{y}|2n,\boldsymbol{p}), \tag{S7}$$

where $1(\cdot)$ is the indicator function. To derive (S5)–(S7), we can consider the following generative process when there is random mating. First, in (S7), $2n$ gamete dosages are randomly drawn from the population using the gamete frequencies $\boldsymbol{p}$. Then, these gametes are randomly paired up, with the distribution of the number of each pairing given a fixed number of categories described in Levene [1949] (their equation (5)), and this results in (S6). Finally, for (S5) we merely calculate $\boldsymbol{x}$ from $\boldsymbol{A}$ using (S2).

To calculate the marginal likelihood under the null, let $\mathcal{T}(\boldsymbol{x})$ be the set of non-negative integer matrices $\boldsymbol{A}$ such that (S2) holds, and let $\boldsymbol{y}(\boldsymbol{A})$ denote the function (S3) that maps $\boldsymbol{A}$ to $\boldsymbol{y}$. Then we have, using prior (7),

$$\Pr(\boldsymbol{x}|H_0) = \int_{\boldsymbol{p}} f(\boldsymbol{x}|n,\boldsymbol{p})\,\text{Dirichlet}(\boldsymbol{p}|\boldsymbol{\alpha}) \tag{S8}$$

$$= \int_{\boldsymbol{p}}\sum_{\boldsymbol{A}}\sum_{\boldsymbol{y}} f(\boldsymbol{x}|\boldsymbol{A})f(\boldsymbol{A}|\boldsymbol{y})f(\boldsymbol{y}|\boldsymbol{p})\,\text{Dirichlet}(\boldsymbol{p}|\boldsymbol{\alpha}) \tag{S9}$$

$$= \sum_{\boldsymbol{A}}\sum_{\boldsymbol{y}} f(\boldsymbol{x}|\boldsymbol{A})f(\boldsymbol{A}|\boldsymbol{y})\,\text{DM}(\boldsymbol{y}|\boldsymbol{\alpha}) \tag{S10}$$

$$= \sum_{\boldsymbol{A}\in\mathcal{T}(\boldsymbol{x})}\sum_{\boldsymbol{y}} f(\boldsymbol{A}|\boldsymbol{y})\,\text{DM}(\boldsymbol{y}|\boldsymbol{\alpha}) \tag{S11}$$

$$= \sum_{\boldsymbol{A}\in\mathcal{T}(\boldsymbol{x})} f(\boldsymbol{A}|\boldsymbol{y}(\boldsymbol{A}))\,\text{DM}(\boldsymbol{y}(\boldsymbol{A})|\boldsymbol{\alpha}). \tag{S12}$$

Line (S12) follows from the previous line because $f(\boldsymbol{A}|\boldsymbol{y})$ is 0 wherever (S3) does not hold. Equation (S12) is a closed form equation for the marginal likelihood under the null.

The summation (S12) turns out to be computationally feasible to calculate for tetraploids ($K = 4$) and for small sample sizes of hexaploids ($K = 6$ and $n \lessapprox 100$). Let's discuss how we sum over $\mathcal{T}(\boldsymbol{x})$. Given $x_k$ we need to sum over all possible tuples in the anti-diagonals of $\boldsymbol{A}$ that correspond to that $x_k$. Let $\boldsymbol{a}_k$ be a vector of $a_{ij}$ elements, with $j \geq i$, such that $i + j = k$ for $k = 0, 1, \ldots, K$, ordered in (reverse reflected) lexicographic order. Note that $\boldsymbol{a}_k$ is the $k$th anti-diagonal of the matrix $\boldsymbol{A}$. For example, consider an octoploid with $K = 8$, then

$$\begin{aligned}&\boldsymbol{a}_0 = (a_{00}),\ \boldsymbol{a}_1 = (a_{01}),\ \boldsymbol{a}_2 = (a_{02}, a_{11}),\ \boldsymbol{a}_3 = (a_{03}, a_{12}),\ \boldsymbol{a}_4 = (a_{04}, a_{13}, a_{22}),\\ &\boldsymbol{a}_5 = (a_{14}, a_{23}),\ \boldsymbol{a}_6 = (a_{24}, a_{33}),\ \boldsymbol{a}_7 = (a_{34}),\ \text{and } \boldsymbol{a}_8 = (a_{44}).\end{aligned} \tag{S13}$$

Each $\boldsymbol{a}_k$ vector is of length $\ell_k := \min\left(\left\lfloor \frac{k}{2} \right\rfloor, \left\lfloor \frac{K-k}{2} \right\rfloor\right) + 1$, according to Lemma S1.

**Lemma S1.** *The number of summands in* (S2) *is*

$$\min\left(\left\lfloor \frac{k}{2} \right\rfloor, \left\lfloor \frac{K-k}{2} \right\rfloor\right) + 1. \tag{S14}$$

*This also corresponds to the number of $a_{ij}$ elements that correspond to a given $x_k$.*

*Proof.* We can re-write the indices of sum (S2) by the following inequalities

$$\begin{aligned}&2i \leq k \text{ and } 0 \leq i \leq K/2 \text{ and } 0 \leq k - i \leq K/2 &\text{(S15)}\\ &\Leftrightarrow i \leq \lfloor k/2 \rfloor \text{ and } 0 \leq i \leq K/2 \text{ and } k - K/2 \leq i \leq k &\text{(S16)}\\ &\Leftrightarrow \max(0, k - K/2) \leq i \leq \lfloor k/2 \rfloor. &\text{(S17)}\end{aligned}$$

Taking the difference of the upper and lower bounds, the number of elements in this sum is $\min[\lfloor k/2 \rfloor, \lfloor (K-k)/2 \rfloor] + 1$. □

Let $\mathcal{S}(\ell_k, x_k)$ be the set of $\ell_k$-tuples that sum to $x_k$. Then we iterate through $\mathcal{T}(\boldsymbol{x})$ by (i) iterating through $k$ and (ii) within each $k$ iterating $\boldsymbol{a}_k$ through $\mathcal{S}(\ell_k, x_k)$. The number of elements in $\mathcal{T}(\boldsymbol{x})$ is provided by Theorem S1.

**Theorem S1.** *Let $\mathcal{T}(\boldsymbol{x})$ be the set of non-negative integer matrices $\boldsymbol{A}$ such that* (S2) *holds. Then, for $K \geq 4$, the number of elements in $\mathcal{T}(\boldsymbol{x})$ is*

$$\prod_{k=2}^{K-2} \binom{x_k + \min\left(\lfloor k/2 \rfloor, \lfloor (K-k)/2 \rfloor\right)}{x_k}. \tag{S18}$$

*Proof.* We need to just take the product of the number of $\boldsymbol{a_k}$ tuples that sum to $\boldsymbol{x}_k$. By Lemma S1, the tuple size corresponding to $x_k$ is $\min\left(\left\lfloor \frac{k}{2} \right\rfloor, \left\lfloor \frac{K-k}{2} \right\rfloor\right) + 1$, and so by classic methods of combinatorics, the number of possible tuples that sum to $x_k$ is

$$\binom{x_k + \min\left(\lfloor k/2 \rfloor, \lfloor (K-k)/2 \rfloor\right)}{x_k}. \tag{S19}$$

When $k \in \{0, 1, K-1, K\}$, this binomial coefficient is just 1, and so taking the product over all $k = 0, 1, ..., K$ results in (S18). □

For $K = 4$ (S18) corresponds to $x_2 + 1$ summands in (S12), and for $K = 6$ this corresponds to $(x_2 + 1)(x_3 + 1)(x_4 + 1)$ summands in (S12). This is computationally feasible for moderate sample sizes. However, for $K = 8$, there are $(x_2 + 1)(x_3 + 1)(x_4 + 2)(x_4 + 1)(x_5 + 1)(x_6 + 1)/2$ summands in (S12), which is computationally infeasible except for the smallest of sample sizes. Therefore, a new strategy is needed.

For larger ploidies, we resort to simulation approaches to obtain the marginal likelihood under the null. We created a Gibbs sampler [Gelfand and Smith, 1990], via the same data augmentation scheme (S5)–(S7), to sample from the posterior under the null. We then used the method of Chib [1995] to estimate the marginal likelihood from this Gibbs sampler. The Gibbs sampler is provided in Procedure S1, and its derivation is provided in Appendix S2.

To summarize the results of this section, we obtain the marginal likelihood under the alternative, $P(\boldsymbol{x}|H_1)$, via (S1). For small ploidies, we obtain the marginal likelihood under the null, $P(\boldsymbol{x}|H_0)$, via (S12). For larger ploidies, we estimate the marginal likelihood under the null via Procedure S1, which uses a data augmentation sampler described in Procedure S2. After obtaining these marginal likelihoods, we obtain the Bayes factor (6).

#### S2 Gibbs sampler using known genotypes

For larger ploidies, we resort to simulation approaches to obtain the marginal likelihood under the null. We will create a Gibbs sampler [Gelfand and Smith, 1990], via the same data augmentation scheme (S5)–(S7), to sample from the posterior under the null. Then we will use the method of Chib [1995] to estimate the marginal likelihood from this Gibbs sampler. Let us begin with the data-augmented Gibbs sampler, represented visually in Figure 1. We can represent the complete likelihood in the following two forms

$$f(\boldsymbol{x}, \boldsymbol{A}|n, \boldsymbol{p}) = f(\boldsymbol{x}|\boldsymbol{A})f(\boldsymbol{A}|\boldsymbol{p}) \tag{S20}$$

$$f(\boldsymbol{x}, \boldsymbol{A}, \boldsymbol{y}|n, \boldsymbol{p}) = f(\boldsymbol{x}|\boldsymbol{A})f(\boldsymbol{A}|\boldsymbol{y})f(\boldsymbol{y}|\boldsymbol{p}). \tag{S21}$$

Both representations (S20) and (S21) will prove useful. All of the terms in (S21) were defined in (S5)–(S7). In (S20), $f(\boldsymbol{A}|\boldsymbol{p})$ is a multinomial distribution

$$(a_{11}, a_{12}, a_{22}, \ldots, a_{K/2,K/2}) \sim \text{Multinom}(n, \boldsymbol{r}), \text{ where } r_{ij} = \begin{cases} p_i^2 & \text{if } i = j, \text{ and} \\ 2p_ip_j & \text{if } i \neq j. \end{cases} \tag{S22}$$

Our Gibbs sampler will iteratively sample $[\boldsymbol{y}|\boldsymbol{x}, \boldsymbol{p}]$ and $[\boldsymbol{p}|\boldsymbol{y}]$. The full conditional of $\boldsymbol{p}$, assuming prior (7), is simply

$$[\boldsymbol{p}|\boldsymbol{y}] \sim \text{Dirichlet}(\boldsymbol{y} + \boldsymbol{\alpha}). \tag{S23}$$

We sample $[\boldsymbol{y}|\boldsymbol{x}, \boldsymbol{p}]$ by first sampling $[\boldsymbol{A}|\boldsymbol{x}, \boldsymbol{p}]$ and then calculating $\boldsymbol{y}$ through (S3). Recall $\boldsymbol{a}_k$ is the $k$th anti-diagonal of $\boldsymbol{A}$. Let $\boldsymbol{c}_k$ be the vector of corresponding $r_{ij}$'s from (S22). For example, in octoploids we have

$$\begin{aligned} &\boldsymbol{c}_0 = (r_{00}),\ \boldsymbol{c}_1 = (r_{01}),\ \boldsymbol{c}_2 = (r_{02}, r_{11}),\ \boldsymbol{c}_3 = (r_{03}, r_{12}),\ \boldsymbol{c}_4 = (r_{04}, r_{13}, r_{22}), \\ &\boldsymbol{c}_5 = (r_{14}, r_{23}),\ \boldsymbol{c}_6 = (r_{24}, r_{33}),\ \boldsymbol{c}_7 = (r_{34}), \text{ and } \boldsymbol{c}_8 = (r_{44}). \end{aligned} \tag{S24}$$

Using (S2) and (S22), we have

$$[\boldsymbol{a}_k|x_k, \boldsymbol{p}] \sim \text{Multinom}(x_k, \boldsymbol{d}_k), \text{ where } d_{k\ell} = \frac{c_{k\ell}}{\sum_\ell c_{k\ell}}. \tag{S25}$$

This will populate the $\boldsymbol{A}$ matrix, from which we can calculate $\boldsymbol{y}$ by (S3). A summary of how to sample $\boldsymbol{y}$ from its full conditional distribution is provided in Procedure S2.

We sample $[\boldsymbol{p}|\boldsymbol{y}]$ and $[\boldsymbol{y}|\boldsymbol{x},\boldsymbol{p}]$ for $B$ iterations, after some initial burn-in period. During this burn-in period, we select $\tilde{\boldsymbol{p}}$ to be the value of $\boldsymbol{p}$ that has the highest posterior probability.

$$\tilde{\boldsymbol{p}} = \underset{\boldsymbol{p} \text{ in burn-in}}{\arg\max} \text{ Multinom}(\boldsymbol{x}|n, \boldsymbol{p} * \boldsymbol{p}) \text{ Dirichlet}(\boldsymbol{p}|\boldsymbol{\alpha}) \tag{S26}$$

Choosing a value of $\tilde{\boldsymbol{p}}$ with high posterior probability will decrease the variance of our marginal likelihood estimate. Let $\boldsymbol{y}^{(b)}$ be the $b$th draw of $\boldsymbol{y}$ from this Gibbs sampler, after burn-in. Then we estimate the posterior evaluated at $\tilde{\boldsymbol{p}}$ by using the method of Chib [1995]

$$\hat{\pi}(\tilde{\boldsymbol{p}}|\boldsymbol{x}) = \frac{1}{B}\sum_{b=1}^{B} \text{Dirichlet}(\tilde{\boldsymbol{p}}|\boldsymbol{y}^{(b)} + \boldsymbol{\alpha}). \tag{S27}$$

By the method of Chib [1995], we estimate the marginal likelihood by

$$\widehat{\Pr}(\boldsymbol{x}|H_0) = \frac{\text{Multinom}(\boldsymbol{x}|n, \tilde{\boldsymbol{p}} * \tilde{\boldsymbol{p}}) \text{ Dirichlet}(\tilde{\boldsymbol{p}}|\boldsymbol{\alpha})}{\hat{\pi}(\tilde{\boldsymbol{p}}|\boldsymbol{x})}. \tag{S28}$$

We summarize the Gibbs sampler and the marginal likelihood estimation approach in Procedure S1.

#### S3 Gibbs sampler using genotype likelihoods

Let $\ell_{ik}$ be the genotype likelihood for individual $i = 1, 2, \ldots, n$ for genotype $k = 0, 1, \ldots, K$. We introduce a new latent variable, $z_i \in \{0, 1, \ldots, K\}$, which is the unobserved genotype for individual $i$. If we represent the data used for genotyping individual $i$ as $d_i$, then $\ell_{ik} = Pr(d_i|z_i = k)$. We have that $z_i$ follows a categorical distribution with support vector $(0, 1, \ldots, K)$ and probability vector $\boldsymbol{q} = (q_0, q_1, \ldots, q_K)$. Under the null, this probability vector can be written as $\boldsymbol{q} = \boldsymbol{p} * \boldsymbol{p}$, where $\boldsymbol{p} = (p_0, p_1, \ldots, p_{K/2})$.

For the Bayes factor, we will need the marginal likelihoods under both the null and the alternative. Calculating the marginal likelihood under the null will again require developing a Gibbs sampler and using the method of Chib [1995]. The joint distribution of data and parameters may be written in the following two useful forms

$$\left[\prod_{i=1}^{n} \Pr(d_i|z_i = k)\right] f(\boldsymbol{z}|\boldsymbol{x}) f(\boldsymbol{x}|\boldsymbol{A}) f(\boldsymbol{A}|\boldsymbol{y}) f(\boldsymbol{y}|\boldsymbol{p}) f(\boldsymbol{p}|\boldsymbol{\alpha}), \text{ and} \tag{S29}$$

$$\left[\prod_{i=1}^{n} \Pr(d_i|z_i = k)\right] f(\boldsymbol{z}|\boldsymbol{p}) f(\boldsymbol{p}|\boldsymbol{\alpha}) \tag{S30}$$

where $\ell_{ik} = \Pr(d_i|z_i = k)$, and $\boldsymbol{z} = (z_1, z_2, \ldots, z_n)$ is found from $\boldsymbol{x}$ by randomly permuting genotypes. The form of $f(\boldsymbol{z}|\boldsymbol{x})$ is not important as $f(\boldsymbol{x}|\boldsymbol{z})$ is merely the indicator that $x_k = \sum_{i=1}^{n} 1(z_i = k)$ is satisfied for each $k$, and this indicator is what is used in the following Gibbs sampler. We have $f(\boldsymbol{z}|\boldsymbol{p}) = \prod_{i=1}^{n} Pr(z_i = k|\boldsymbol{p})$, where $Pr(z_i = k|\boldsymbol{p})$ is the $k$th element of $\boldsymbol{p} * \boldsymbol{p}$. The form for $f(\boldsymbol{x}|\boldsymbol{A})$, $f(\boldsymbol{y}|\boldsymbol{p})$ and $f(\boldsymbol{p}|\boldsymbol{\alpha})$ are the same as in Section S1. The Gibbs sampler consists of iteratively doing the following

1. Sample $[\boldsymbol{z}|\boldsymbol{p},\boldsymbol{d}]$ by the following two steps:
   (a) Calculate $\boldsymbol{q}=\boldsymbol{p}*\boldsymbol{p}$.
   (b) Sample $z_i=k$ with probability $\ell_{ik}q_k/\sum_{k=0}^K \ell_{ik}q_k$.
2. Calculate $x_k=\sum_{i=1}^n 1(z_i=k)$.
3. Sample $[\boldsymbol{y}|\boldsymbol{x},\boldsymbol{p}]$ using Procedure S2.
4. Sample $[\boldsymbol{p}|\boldsymbol{y}]\sim \text{Dirichlet}(\boldsymbol{y}+\boldsymbol{\alpha})$.

We now describe how to use the method of Chib [1995] to obtain the marginal likelihood under the null. First, we note that

$$f(\boldsymbol{d}|\tilde{\boldsymbol{p}})=\prod_{i=1}^n\sum_{k=0}^K \ell_{ik}\tilde{q}_k \text{ where } \tilde{\boldsymbol{q}}=\tilde{\boldsymbol{p}}*\tilde{\boldsymbol{p}}, \text{ and} \tag{S31}$$

$$f(\tilde{\boldsymbol{p}})=\text{Dirichlet}(\tilde{\boldsymbol{p}}|\boldsymbol{\alpha}). \tag{S32}$$

To obtain $\pi(\tilde{\boldsymbol{p}}|\boldsymbol{d})$, we let $\boldsymbol{y}^{(b)}$ be the $b$th sample of $\boldsymbol{y}$ from the Gibbs sampler, which marginally follows the distribution $f(\boldsymbol{y}|\boldsymbol{d})$. Since

$$\pi(\tilde{\boldsymbol{p}}|\boldsymbol{d})=\int_{\boldsymbol{y}} f(\boldsymbol{p}|\boldsymbol{y},\boldsymbol{d})f(\boldsymbol{y}|\boldsymbol{d})\mathrm{d}\boldsymbol{y}=\int_{\boldsymbol{y}} f(\boldsymbol{p}|\boldsymbol{y})f(\boldsymbol{y}|\boldsymbol{d})\mathrm{d}\boldsymbol{y}, \tag{S33}$$

we estimate the posterior by

$$\hat{\pi}(\tilde{\boldsymbol{p}}|\boldsymbol{d})=\frac{1}{B}\sum_{b=1}^B \text{Dirichlet}(\tilde{\boldsymbol{p}}|\boldsymbol{y}^{(b)}+\boldsymbol{\alpha}). \tag{S34}$$

The marginal likelihood is then estimated by

$$\widehat{\Pr}(\boldsymbol{d}|H_0)=\frac{\text{Dirichlet}(\tilde{\boldsymbol{p}}|\boldsymbol{\alpha})\prod_{i=1}^n\sum_{k=0}^K \ell_{ik}\tilde{q}_k}{\hat{\pi}(\tilde{\boldsymbol{p}}|\boldsymbol{d})}. \tag{S35}$$

The choice of $\tilde{\boldsymbol{p}}$ is only important in that values with high probability result in a lower variance of the estimated marginal likelihood. We again choose the $\tilde{\boldsymbol{p}}$ that yields the highest posterior probability during the burn-in period.

$$\tilde{\boldsymbol{p}}=\underset{\boldsymbol{p}\text{ in burn-in}}{\arg\max}\ \text{Dirichlet}(\boldsymbol{p}|\boldsymbol{\alpha})\prod_{i=1}^n\sum_{k=0}^K \ell_{ik}q_k, \text{ where } \boldsymbol{q}=\boldsymbol{p}*\boldsymbol{p}. \tag{S36}$$

Our approach to estimate the marginal likelihood under the null, using genotype likelihoods, is summarized in Procedure S3.

Let us turn to the marginal likelihood under the alternative where, unlike in Section S1, we now need to implement a Gibbs sampler to calculate it. We again suppose that *a prior* $\boldsymbol{q}$ follows a Dirichlet($\boldsymbol{\beta}$) distribution. Then we can write the joint distribution of the data and parameters as

$$\left[\prod_{i=1}^n \Pr(d_i|z_i=k)\right] f(\boldsymbol{z}|\boldsymbol{q})f(\boldsymbol{q}|\boldsymbol{\beta}), \tag{S37}$$

where $\ell_{ik} = \Pr(d_i|z_i = k)$, $f(\boldsymbol{z}|\boldsymbol{p}) = \prod_{i=1}^n Pr(z_i = k|\boldsymbol{q})$ where $Pr(z_i = k|\boldsymbol{q}) = q_k$, and $f(\boldsymbol{q}|\boldsymbol{\beta})$ is the probability mass function of the Dirichlet distribution with parameter $\boldsymbol{\beta}$ evaluated at $\boldsymbol{q}$. The Gibbs sampler consists of iteratively performing the following steps:

1. Sample $[\boldsymbol{z}|\boldsymbol{q}]$: sample $z_i = k$ with probability $\ell_{ik}q_k / \sum_{k=0}^K \ell_{ik}q_k$.
2. Calculate $x_k = \sum_{i=1}^n 1(z_i = k)$
3. Sample $[\boldsymbol{q}|\boldsymbol{z}]$ from Dirichlet$(\boldsymbol{x} + \boldsymbol{\beta})$

To obtain $\hat{\pi}(\boldsymbol{q}|\boldsymbol{d})$, we let $\boldsymbol{x}^{(b)}$ be the $b$th value of $\boldsymbol{x}$. Then we estimate the posterior density at some value $\tilde{\boldsymbol{q}}$ by

$$\hat{\pi}(\tilde{\boldsymbol{q}}|\boldsymbol{d}) = \frac{1}{B}\sum_{b=1}^B \text{Dirichlet}(\tilde{\boldsymbol{q}}|\boldsymbol{x}^{(b)} + \boldsymbol{\beta}). \tag{S38}$$

The marginal likelihood is estimated, using the method of Chib [1995], by

$$\widehat{\Pr}(\boldsymbol{d}|H_1) = \frac{\text{Dirichlet}(\tilde{\boldsymbol{q}}|\boldsymbol{\beta})\prod_{i=1}^n \sum_{k=0}^K \ell_{ik}\tilde{q}_k}{\hat{\pi}(\tilde{\boldsymbol{q}}|\boldsymbol{d})}. \tag{S39}$$

Here, we choose a value of $\tilde{\boldsymbol{q}}$ that has large posterior probability. Specifically, we choose

$$\tilde{\boldsymbol{q}} = \underset{\boldsymbol{q} \text{ in burn-in}}{\arg\max}\ \text{Dirichlet}(\boldsymbol{q}|\boldsymbol{\beta})\prod_{i=1}^n\sum_{k=0}^K \ell_{ik}q_k. \tag{S40}$$

### S4 Proof for prior elicitation of $\boldsymbol{\beta}$

To prove result (22), first note that under the alternative we have $\boldsymbol{x} \sim \text{DM}(1, \boldsymbol{\beta})$. Under a sample of size 1, the Dirichlet-Multinomial reduces to a categorical distribution with $\Pr(\boldsymbol{x} = \boldsymbol{e}_k) = \beta_k / \sum_{k=0}^K \beta_k$. Thus, the scaling of $\boldsymbol{\beta}$ does not impact property (21). This means that we just need to find $\Pr(\boldsymbol{x} = \boldsymbol{e}_k)$ under the null and set the $\beta_k$'s accordingly to match these probabilities. This will result in satisfying property (21).

**Theorem S2.** *Let $\mathcal{T}(\boldsymbol{x})$ be the space of non-negative integer matrices $\boldsymbol{A}$ that satisfy (S2). Let $\boldsymbol{y}(\boldsymbol{A})$ be the function that maps $\boldsymbol{A}$ to $\boldsymbol{y}$ via (S3). Then setting*

$$\beta_k \propto \sum_{\boldsymbol{A}\in\mathcal{T}(\boldsymbol{e}_k)} \text{DM}(\boldsymbol{y}(\boldsymbol{A})|\boldsymbol{\alpha}) \tag{S41}$$

*will result in property (21).*

*Proof.* We just need to calculate $f(\boldsymbol{e}_k)$ for each $k = 0, 1, \ldots, K$. Since marginally, *a priori*, $\boldsymbol{y} \sim \text{DM}(\boldsymbol{\alpha})$, we have

$$f(\boldsymbol{e}_k) = \sum_{\boldsymbol{A}}\sum_{\boldsymbol{y}} f(\boldsymbol{e}_k|\boldsymbol{A})f(\boldsymbol{A}|\boldsymbol{y})\,\text{DM}(\boldsymbol{y}|\boldsymbol{\alpha}), \tag{S42}$$

where we recall that $f(\boldsymbol{e}_k|\boldsymbol{A})$ is the indicator that (S2) holds, and $f(\boldsymbol{A}|\boldsymbol{y})$ is from (S6).

By (S3) each $\boldsymbol{A}$ corresponds to a single $\boldsymbol{y}$. The main trick of this proof is to note that, when $n = 1$, we also have that each $\boldsymbol{y}$ corresponds to a single $\boldsymbol{A}$ by

$$a_{ij} = \begin{cases} 1 & \text{if } i \neq j \text{ and } y_i = y_j = 1, \\ 1 & \text{if } i = j \text{ and } y_i = 2, \\ 0 & \text{otherwise.} \end{cases} \tag{S43}$$

That means that $f(\boldsymbol{A}|\boldsymbol{y})$ is now just the indicator that (S43) holds. That means we can re-write (S42) as

$$\begin{aligned} f(\boldsymbol{e}_k) &= \sum_{\boldsymbol{A}} \sum_{\boldsymbol{y}} f(\boldsymbol{e}_k|\boldsymbol{A}) f(\boldsymbol{A}|\boldsymbol{y}) \, \mathrm{DM}(\boldsymbol{y}|\boldsymbol{\alpha}) && \text{(S44)} \\ &= \sum_{\boldsymbol{A} \in \mathcal{T}(\boldsymbol{e}_k)} \sum_{\boldsymbol{y}} f(\boldsymbol{A}|\boldsymbol{y}) \, \mathrm{DM}(\boldsymbol{y}|\boldsymbol{\alpha}) && \text{(S45)} \\ &= \sum_{\boldsymbol{A} \in \mathcal{T}(\boldsymbol{e}_k)} \mathrm{DM}(\boldsymbol{y}(\boldsymbol{A})|\boldsymbol{\alpha}). && \text{(S46)} \end{aligned}$$

□

For the special case when $\boldsymbol{\alpha} = \mathbf{1}_{K/2+1}$, we will need Lemma S2.

**Lemma S2.** *Suppose* $\boldsymbol{p} \sim \mathrm{Dirichlet}(\mathbf{1}_{K/2+1})$ *and* $[\boldsymbol{y}|\boldsymbol{p}] \sim \mathrm{Multinom}(2n, \boldsymbol{p})$. *Then*

$$\Pr(\boldsymbol{y}) = \frac{1}{\binom{K/2+2n}{K/2}}. \tag{S47}$$

*That is, marginally,* $\boldsymbol{y}$ *is discrete uniform over the space of* $(K/2+1)$*-tuples of non-negative integers whose sum is* $2n$.

*Proof.* Marginally, we have $\boldsymbol{y} \sim \mathrm{DM}(2n, \mathbf{1}_{K/2+1})$. Calculating the density of the Dirichlet-multinomial, we have

$$\begin{aligned} \Pr(\boldsymbol{y}) &= \frac{\Gamma(K/2+1)\Gamma(2n+1)}{\Gamma(2n+K/2+1)} \prod_{k=0}^{K} \frac{\Gamma(y_k+1)}{\Gamma(1)\Gamma(y_k+1)} && \text{(S48)} \\ &= \frac{(K/2)!(2n)!}{(K/2+2n)!} && \text{(S49)} \\ &= \frac{1}{\binom{K/2+2n}{K/2}}. && \text{(S50)} \end{aligned}$$

□

We then have Corollary S1.

**Corollary S1.** *Suppose* $\boldsymbol{p} \sim \mathrm{Dirichlet}(\mathbf{1}_{K/2+1})$ *and* $[\boldsymbol{x}|\boldsymbol{p}] \sim \mathrm{Multinom}(1, \boldsymbol{p} * \boldsymbol{p})$. *Then setting*

$$\beta_k \propto \frac{\min\left(\left\lfloor \frac{k}{2} \right\rfloor, \left\lfloor \frac{K-k}{2} \right\rfloor\right) + 1}{\binom{K/2+2}{K/2}} \tag{S51}$$

*will result in property* (21).

*Proof.* From Theorem S2, we just need to calculate $\sum_{\boldsymbol{A}\in\mathcal{T}(\boldsymbol{e}_k)} \mathrm{DM}(\boldsymbol{y}(\boldsymbol{A})|\boldsymbol{1}_{K/2+1})$. But by Lemma S2, $\boldsymbol{y}$ is uniformly distributed, so $\mathrm{DM}(\boldsymbol{y}(\boldsymbol{A})|\boldsymbol{1}_{K/2+1})$ is the same for all $\boldsymbol{y}$. Using Lemma S1, there are $\min[\lfloor k/2\rfloor, \lfloor (K-k)/2\rfloor]+1$ elements in this summation (the number of ways to place the single non-zero element in the matrix $\boldsymbol{A}$). Thus, we have

$$f(\boldsymbol{e}_k) = \frac{\min\left(\left\lfloor \frac{k}{2}\right\rfloor, \left\lfloor \frac{K-k}{2}\right\rfloor\right) + 1}{\binom{K/2+2}{K/2}}. \tag{S52}$$

□

##### S4.1 Default priors for allopolyploids

Under HWP, the alleles of an allopolyploid are just a random sample from each subgenome's pool, where the probability of randomly sampling the minor allele from subgenome $j$ is, say, $r_j$ (subgenome $j$'s allele frequency). Then, we have that

$$p_k = \sum_{\substack{\boldsymbol{z}\in\{0,1\}^{K/2} \\ \text{s.t.} \\ \sum_{j=1}^{K/2} z_j = k}} \prod_{j=1}^{K/2} (1-r_j)^{1-z_j} r_j^{z_j}. \tag{S53}$$

Equation (S53) is merely summing over the number of ways to obtain a gamete dosage of $k$ from $K/2$ subgenomes where the probability of a minor allele is $r_j$ for subgenome $j$. It is natural to place a uniform prior over $r_j$, and this induces a prior distribution over $\boldsymbol{p}$. We can moment match with the Dirichlet($\boldsymbol{\alpha}$) distribution like in Section 2.3 to obtain $\boldsymbol{\alpha}$. Doing this, we have

$$\alpha_k \propto \mathrm{E}[p_k] \tag{S54}$$

$$= \sum_{\substack{\boldsymbol{z}\in\{0,1\}^{K/2} \\ \text{s.t.} \\ \sum_{j=1}^{K/2} z_j = k}} \prod_{j=1}^{K/2} \mathrm{E}\left[(1-r_j)^{1-z_j} r_j^{z_j}\right] \tag{S55}$$

$$= \sum_{\substack{\boldsymbol{z}\in\{0,1\}^{K/2} \\ \text{s.t.} \\ \sum_{j=1}^{K/2} z_j = k}} \prod_{j=1}^{K/2} \frac{1}{2} \tag{S56}$$

$$= \frac{1}{2^{K/2}}\binom{K/2}{k}, \tag{S57}$$

where line (S56) follows from symmetry (both $r_j$ and $1-r_j$ follow a Unif$[0,1]$, and each $z_j$ is either 0 or 1, so $\mathrm{E}[(1-r_j)^{1-z_j} r_j^{z_j}] = 1/2$). Note that (S57) are just binomial frequencies. By default, we use

$$\alpha_k = \binom{K/2}{k}\frac{K+2}{2^{K/2+1}}, \tag{S58}$$

so that the total sum of $\boldsymbol{\alpha}$ is the same as $\mathbf{1}_{K/2+1}$. For $\boldsymbol{\beta}$, we use the same strategy as in Section 2.3 to obtain a $\boldsymbol{\beta}$ that results in a Bayes factor of 1 for all samples of size 1 (Appendix S4).

#### S5 Prior sensitivity analysis

Bayes factors are known to be sensitive to prior selection, even for large sample sizes [Kass and Raftery, 1995]. Thus, to explore the impact of our prior selection, we included in the simulations of Section 3.1 additional choices for $\boldsymbol{\alpha}$. Specifically, we chose

- $\boldsymbol{\alpha}_{\text{auto}} = \mathbf{1}_{K/2+1}$,
- $\boldsymbol{\alpha}_{\text{allo}} = \frac{K+2}{2^{K/2+1}}\left(\binom{K/2}{0}, \binom{K/2}{1}, \ldots, \binom{K/2}{K/2}\right)$,
- $\boldsymbol{\alpha}_1 = \frac{1}{2}\mathbf{1}_{K/2+1}$,
- $\boldsymbol{\alpha}_2 = 2\mathbf{1}_{K/2+1}$,
- $\boldsymbol{\alpha}_3 = \frac{4}{K+4}(1, 2, \ldots, K/2+1)$,

The value of $\boldsymbol{\alpha}_{\text{auto}}$ is our default for autopolyploids, the value of $\boldsymbol{\alpha}_{\text{allo}}$ is our default for allopolyploids, $\boldsymbol{\alpha}_1$ concentrates the prior toward the corners of the simplex, $\boldsymbol{\alpha}_2$ concentrates the prior toward the center of the simplex, and $\boldsymbol{\alpha}_3$ places non-uniform probability on different genotypes but is scaled to have the same weight as $\boldsymbol{\alpha}_{\text{auto}}$. All of these are moderate deviations from the default priors. For each value of $\boldsymbol{\alpha}$, we chose $\boldsymbol{\beta}$ according to the strategy in Section 2.3.

The results are presented in Figure S4. There, we see that any of the five priors explored resulted in consistent behavior of the Bayes factor — increasing with sample size when the null is true and decreasing with sample size when the alternative is true. We see that for moderate deviations from our default priors, the Bayes factors are relatively stable. This indicates that our methods are relatively robust to prior choice.

#### S6 Small sample behavior

We can gain a sense of how our Bayes procedure works on small samples by applying it to all possible values of $\boldsymbol{x}$ when $n$ is small enough for this to be feasible.

We applied the Bayes procedure from Section 2.1 to all $(K+1)$-tuples ($K \in \{4, 6, 8\}$) that sum to $n \in \{5, 10\}$. This corresponds to $\binom{K+n}{K}$ tuples for a given ploidy and sample size. Each tuple represents a possible dataset $\boldsymbol{x}$ and each $\boldsymbol{x}$ has a Bayes factor with a given $\boldsymbol{\alpha}$ and $\boldsymbol{\beta}$. The histogram of log Bayes factors is presented in Figure S2. We see there that for samples of size 5 the log Bayes factors are almost all between -2 and 2, and for samples of size 10 the log Bayes factors are almost all between -7 and 3. The ploidy of the species does not considerably change the distribution of log Bayes factors. Using the suggested thresholds of Bayes factors from Kass and Raftery [1995], this would indicate that it is possible to obtain "very strong" evidence against the null hypothesis at samples as small as size 10, but not at size 5.

In Table S2, we list the values of $\boldsymbol{x}$ that produce the largest and smallest Bayes factors. We see there that the largest Bayes factors tend to result from very unimodal data, whereas the smallest Bayes factors tend to result from data with a couple of prominent modes. This is verified in Figure S3 where we see that unimodal $\boldsymbol{x}$ tends to result in positive log Bayes factors, while non-unimodal $\boldsymbol{x}$ tends to result in negative log Bayes factors. Most, but not all, of the information in the consistency with a random mating hypothesis appears to be in the form of unimodality.

#### S7 A Bayesian test for F1/S1 segregation frequencies

Here, we describe a Bayesian procedure to test for Mendelian segregation frequencies among F1 and S1 populations using genotype likelihoods. Let $\boldsymbol{p}_1$ be the gamete frequencies for parent 1 and let $\boldsymbol{p}_2$ be the gamete frequencies for parent 2, where in an S1 population $\boldsymbol{p}_1 = \boldsymbol{p}_2$. Let $z_j \in \{0, 1, \ldots, K\}$ be the genotype for parent $j \in \{1, 2\}$. Then, under Mendelian segregation, in the absence of double reduction [Huang et al., 2019], we have that the $\boldsymbol{p}_j$'s are Hypergeometric probabilities [Serang et al., 2012]. Specifically,

$$p_{jk} = \mathrm{HG}(k, K/2|z_j, K) = \frac{\binom{z_j}{k}\binom{K-z_j}{K/2-k}}{\binom{K}{K/2}}. \tag{S59}$$

Our test, then is to compare

$$H_0 : \boldsymbol{q} = \boldsymbol{p}_1 * \boldsymbol{p}_2 \text{ where the } \boldsymbol{p}_j\text{'s are from (S59)}, \tag{S60}$$

$$H_1 : \boldsymbol{q} \neq \boldsymbol{p}_1 * \boldsymbol{p}_2. \tag{S61}$$

To implement this test, we need the marginal likelihoods under both the null and alternative. As before, let $\ell_{ik}$ be the genotype likelihood for individual $i$ at genotype $k$. Let $\tilde{\ell}_{jk}$ be the genotype likelihood for parent $j \in \{1, 2\}$ at genotype $k$. The likelihood for the offspring is the same as before (17). For the null model, we just need a prior over the $z_j$'s. We derive these from the genotype likelihoods for the parents

$$\Pr(z_j) = \frac{\tilde{\ell}_{jk}}{\sum_{k=0}^{K} \tilde{\ell}_{jk}}. \tag{S62}$$

If parental data are not available, then we set $\Pr(z_j) = 1/(K + 1)$. We can then calculate the marginal likelihood under the null in closed form,

$$\Pr(\text{data}|H_0) = \sum_{z_1=0}^{K}\sum_{z_2=0}^{K}\left(\frac{\tilde{\ell}_{1k}}{\sum_{k=0}^{K}\tilde{\ell}_{1k}}\right)\left(\frac{\tilde{\ell}_{2k}}{\sum_{k=0}^{K}\tilde{\ell}_{2k}}\right)\prod_{i=1}^{n}\sum_{k=0}^{K}\ell_{ik}\boldsymbol{q}(z_1, z_2)_k, \tag{S63}$$

where $\boldsymbol{q}(z_1, z_2) = \boldsymbol{p}_1 * \boldsymbol{p}_2$ where $\boldsymbol{p}_j$ is defined in (S59).

For the alternative model, using prior (8), we use the same Stan [Stan Development Team, 2022a,b] implementation as in Section 2.2, and use bridge sampling [Meng and Wong, 1996, Gronau et al., 2020] to estimate the marginal likelihood. After obtaining both marginal likelihoods, we can calculate the Bayes factor.

#### S8 Random mating versus Hardy-Weinberg proportions in allopolyploids

In this section, we detail how random mating and HWP differ in allopolyploids, describe how to update genotype frequencies in an allotetraploid population after a generation of random mating, and demonstrate empirically how HWP are only approached asymptotically in an allotetraploid population.

The Hardy-Weinberg proportions for allopolyploids occur if (i) each subgenome separately has diploid HWP and (ii) each subgenome is independent. If conditions (i) and (ii) are both met, then the HWP for an allopolyploid population of ploidy $K$ can be represented by

$$\boldsymbol{q} = \boldsymbol{s}_1 * \boldsymbol{s}_2 * \cdots * \boldsymbol{s}_{K/2}, \tag{S64}$$

where $\boldsymbol{s}_k = (1-r_k, r_k) * (1-r_k, r_k) = [(1-r_k)^2, 2r_k(1-r_k), r_k^2]$ are the Hardy-Weinberg proportions of subgenome $k$ with allele frequency $r_k$. For example, in an allotetraploid the HWP are

$$q_0 = (1 - r_1)^2(1 - r_2)^2, \tag{S65}$$
$$q_1 = 2(1 - r_1)^2 r_2(1 - r_2) + 2r_1(1 - r_1)(1 - r_2)^2, \tag{S66}$$
$$q_2 = (1 - r_1)^2 r_2^2 + 4r_1(1 - r_1)r_2(1 - r_2) + r_1^2(1 - r_2)^2, \tag{S67}$$
$$q_3 = 2r_1(1 - r_1)r_2^2 + 2r_1^2 r_2(1 - r_2), \text{ and} \tag{S68}$$
$$q_4 = r_1^2 r_2^2. \tag{S69}$$

Condition (i) will occur after one or two generations of random mating. That is, each subgenome will separately have HWP after one or two generations. However, the dependence between subgenomes for condition (ii) may take many rounds of random mating to dissipate. That is, if there is any dependence between the subgenomes, HWP in allopolyploids is also only approached asymptotically.

Let us describe the next generation's genotype frequencies under random mating in allotetraploids. Let $\boldsymbol{G} = (g_{ij}) \in \mathbb{R}^{3\times 3}$ be a matrix where $g_{ij}$ is the frequency of allotetraploid individuals with genotypes $i \in \{0, 1, 2\}$ in the first subgenome and $j \in \{0, 1, 2\}$ in the second subgenome. The genotype frequencies are the sums of the anti-diagonals of $\boldsymbol{G}$,

$$q_k = \sum_i g_{i,k-i}. \tag{S70}$$

The subgenome genotype frequencies are the row and column sums of $\boldsymbol{G}$. The gamete frequencies of this population are

$$\boldsymbol{P} = \begin{pmatrix} g_{00} + g_{01}/2 + g_{10}/2 + g_{11}/4 & g_{01}/2 + g_{02} + g_{11}/4 + g_{12}/2 \\ g_{10}/2 + g_{11}/4 + g_{20} + g_{21}/4 & g_{11}/4 + g_{12}/2 + g_{21}/2 + g_{22} \end{pmatrix}, \tag{S71}$$

where $p_{ij}$ is the proportion of gametes with genotype $i \in \{0, 1\}$ in the first subgenome and $j \in \{0, 1\}$ in the second subgenome. The next generation's joint distribution of subgenome genotypes, under random mating, is

$$\boldsymbol{G}^{\text{next}} = \boldsymbol{P} \overset{2}{*} \boldsymbol{P}, \tag{S72}$$

where $\overset{2}{*}$ is 2-dimensional convolution (S75).

We now empirically demonstrate the asymptotic nature of HWP in allotetraploids. We iteratively applied (S71) and (S72) for 10 generations, starting at a value of $\boldsymbol{G}$ that has moderate dependence between subgenomes,

$$\boldsymbol{G}^{\text{init}} = \begin{pmatrix} .1 & .2 & 0 \\ 0 & .3 & 0 \\ 0 & .1 & .3 \end{pmatrix}. \tag{S73}$$

After the first generation of random mating, the subgenomes both reach HWP. However, HWP for the genotype frequencies (S70) are only approached asymptotically (Figure S23).

### S9 Random mating at multiallelic loci

Here, we detail here how to mathematically represent the hypothesis of random mating at a multi-allelic locus, and discuss how one would approach testing for random mating. Consider a locus with $m+1$ alleles, $A_0, A_1, \ldots, A_m$, in a $K$-ploid species. We collect the gamete frequencies for alleles $A_1, A_2, \ldots, A_m$ into a $m$-dimensional array $\mathcal{P} = (p_{\boldsymbol{k}}) \in \mathbb{R}^{(K/2+1)\times(K/2+1)\times\cdots\times(K/2+1)}$, where for the multi-index $\boldsymbol{k} = (k_1, k_2, \ldots, k_m)$, $p_{\boldsymbol{k}}$ is the proportion of gametes with $k_1$ copies of $A_1$, $k_2$ copies of $A_2$, ..., $k_m$ copies of $A_m$, and, thus, $K/2 - \sum_{j=1}^m k_j$ copies of $A_0$. Each $k_j$ ranges from $0, 1, \ldots K/2$. Because each gamete has $K/2$ genome copies, this implies that $p_{\boldsymbol{k}} = 0$ when $\sum_{j=1}^m k_j > K/2$. Let $\mathcal{Q} = (q_{\boldsymbol{k}}) \in \mathbb{R}^{(K+1)\times(K+1)\times\cdots(K+1)}$ be the $m$-dimensional array of genotype frequencies, defined similarly as $\mathcal{P}$ but with each $k_j$ ranging from $0, 1, \ldots, K$. Then random mating implies that

$$\mathcal{Q} = \mathcal{P} \overset{m}{*} \mathcal{P}, \text{ or} \tag{S74}$$

$$q_{\boldsymbol{k}} = \sum_{\boldsymbol{i}} p_{\boldsymbol{i}} p_{\boldsymbol{k}-\boldsymbol{i}} \tag{S75}$$

where $\overset{m}{*}$ denotes $m$-dimensional discrete linear convolution [Rakhuba and Oseledets, 2015], represented in (S75) where the multi-index $\boldsymbol{i}$ sums over $0 \leq i_1, i_2, \ldots, i_m \leq K/2$. For example, suppose we have a tri-allelic locus in a tetraploid population, then

$$\mathcal{P} = \begin{pmatrix} p_{00} & p_{01} & p_{02} \\ p_{10} & p_{11} & 0 \\ p_{20} & 0 & 0 \end{pmatrix}, \tag{S76}$$

where $p_{ij}$ is the frequency of gametes with $i$ copies of $A_1$, $j$ copies of $A_2$, and $2-i-j$ copies of $A_0$. Under random mating, we have

$$\mathcal{Q} = \mathcal{P} \overset{m}{*} \mathcal{P} = \begin{pmatrix} p_{00}^2 & 2p_{00}p_{01} & p_{01}^2 + 2p_{00}p_{02} & 2p_{01}p_{02} & p_{02}^2 \\ 2p_{00}p_{10} & 2p_{00}p_{11} + 2p_{01}p_{10} & 2p_{01}p_{11} + 2p_{02}p_{10} & 2p_{02}p_{11} & 0 \\ p_{10}^2 + 2p_{00}p_{20} & 2p_{10}p_{11} + 2p_{01}p_{20} & p_{11}^2 + 2p_{02}p_{20} & 0 & 0 \\ 2p_{10}p_{20} & 2p_{11}p_{20} & 0 & 0 & 0 \\ p_{20}^2 & 0 & 0 & 0 & 0 \end{pmatrix}. \tag{S77}$$

Clearly, testing (S74) is much more complicated than testing (3). It might be easiest to avoid a data augmentation scheme like in Section 2.1 and implement a test for (S74) using Stan.

### S10 Supplementary tables, figures, and procedures

**Procedure S1** Gibbs sampler and marginal likelihood estimation under the null of random mating when genotypes are known.

**Input:** Genotype counts $\boldsymbol{x} = (x_0, x_1, \ldots, x_K)$ for a $K$-ploid population, hyperparameter $\boldsymbol{\alpha} = (\alpha_0, \alpha_1, \ldots, \alpha_{K/2})$, $B$ sampling iterations, and $T$ burn-in iterations.
Initialize $\boldsymbol{p} \sim \text{Dirichlet}(\boldsymbol{\alpha})$
Initialize $\tilde{\boldsymbol{p}} \leftarrow \boldsymbol{p}$
Initialize $\tilde{L} \leftarrow \text{Multinom}(\boldsymbol{x}|n, \boldsymbol{p} * \boldsymbol{p})\,\text{Dirichlet}(\boldsymbol{p}|\boldsymbol{\alpha})$
Initialize $\hat{\pi} \leftarrow 0$
**for** $j = 1, 2, \ldots, T + B$ **do**
    Sample $[\boldsymbol{y}|\boldsymbol{x}, \boldsymbol{p}]$ using Procedure S2
    Sample $[\boldsymbol{p}|\boldsymbol{y}] \sim \text{Dirichlet}(\boldsymbol{y} + \boldsymbol{\alpha})$
    **if** $(\text{Multinom}(\boldsymbol{x}|n, \boldsymbol{p} * \boldsymbol{p})\,\text{Dirichlet}(\boldsymbol{p}|\boldsymbol{\alpha}) > \tilde{L})$ **and** $(j \leq T)$ **then**
        $\tilde{\boldsymbol{p}} \leftarrow \boldsymbol{p}$
        $\tilde{L} \leftarrow \text{Multinom}(\boldsymbol{x}|n, \boldsymbol{p} * \boldsymbol{p})\,\text{Dirichlet}(\boldsymbol{p}|\boldsymbol{\alpha})$
    **else if** $j > T$ **then**
        $\hat{\pi} \leftarrow \hat{\pi} + \frac{1}{B}\,\text{Dirichlet}(\tilde{\boldsymbol{p}}|\boldsymbol{y} + \boldsymbol{\alpha})$
    **end if**
**end for**
Estimate marginal likelihood

$$\widehat{\Pr}(\boldsymbol{x}|H_0) = \frac{\text{Multinom}(\boldsymbol{x}|n, \tilde{\boldsymbol{p}} * \tilde{\boldsymbol{p}})\,\text{Dirichlet}(\tilde{\boldsymbol{p}}|\boldsymbol{\alpha})}{\hat{\pi}} \tag{S78}$$

**Return:** Estimate of marginal likelihood $\widehat{\Pr}(\boldsymbol{x}|H_0)$.

| Ploidy | $c$ | $\tilde{\beta}_0$ | $\tilde{\beta}_1$ | $\tilde{\beta}_2$ | $\tilde{\beta}_3$ | $\tilde{\beta}_4$ | $\tilde{\beta}_5$ | $\tilde{\beta}_6$ | $\tilde{\beta}_7$ | $\tilde{\beta}_8$ | $\tilde{\beta}_9$ | $\tilde{\beta}_{10}$ | $\tilde{\beta}_{11}$ | $\tilde{\beta}_{12}$ |
|---|---|---|---|---|---|---|---|---|---|---|---|---|---|---|
| 2 | 1 | 1 | 1 | 1 | - | - | - | - | - | - | - | - | - | - |
| 4 | 5/6 | 1 | 1 | 2 | 1 | 1 | - | - | - | - | - | - | - | - |
| 6 | 7/10 | 1 | 1 | 2 | 2 | 2 | 1 | 1 | - | - | - | - | - | - |
| 8 | 9/15 | 1 | 1 | 2 | 2 | 3 | 2 | 2 | 1 | 1 | - | - | - | - |
| 10 | 11/21 | 1 | 1 | 2 | 2 | 3 | 3 | 3 | 2 | 2 | 1 | 1 | - | - |
| 12 | 13/28 | 1 | 1 | 2 | 2 | 3 | 3 | 4 | 3 | 3 | 2 | 2 | 1 | 1 |

Table S1: Default values for $\beta_k = c\tilde{\beta}_k$, listed for ploidies 2, 4, 6, 8, 10, and 12. The default value for $\boldsymbol{\alpha}$ is $\mathbf{1}_{K/2+1}$.

**Procedure S2** Sampling from the full conditional of $[\boldsymbol{y}|\boldsymbol{x}, \boldsymbol{p}]$.

**Input:** Genotype counts $\boldsymbol{x} = (x_0, x_1, \ldots, x_K)$ for a $K$-ploid population, and gamete frequencies $\boldsymbol{p} = (p_0, p_1, \ldots, p_{K/2})$.

Calculate $r_{ij}$ for $0 \leq i \leq j \leq K/2$:

$$r_{ij} = \begin{cases} p_i^2 & \text{if } i = j, \text{ and} \\ 2p_ip_j & \text{if } i \neq j. \end{cases} \tag{S79}$$

**for** $k = 0, 1, \ldots, K$ **do**

  $\boldsymbol{c}_k = (r_{ij})_{0 \leq i \leq j \leq K/2 \text{ s.t. } i+j=k}$

  $\boldsymbol{d}_{k\ell} = c_{k\ell} / \sum_\ell c_{k\ell}$

  Sample $\boldsymbol{a}_k = (a_{ij})_{0 \leq i \leq j \leq K/2 \text{ s.t. } i+j=k}$ from Multinom$(x_k, \boldsymbol{d}_k)$

**end for**

$\boldsymbol{A}$ is now populated.

**for** $k = 0, 1, \ldots, K/2$ **do**

  $y_k = \sum_{i \text{ s.t. } i<k} a_{ik} + 2a_{kk} + \sum_{j \text{ s.t. } j>k} a_{kj}$.

**end for**

**Return:** Sampled gamete counts $\boldsymbol{y}$.

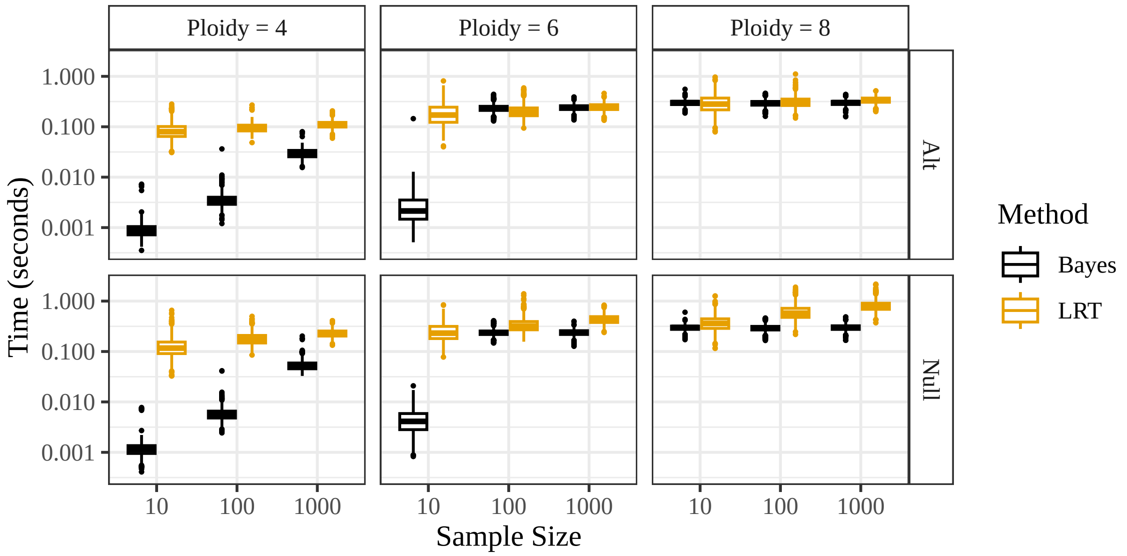


Figure S1: Run time ($y$-axis) for the likelihood ratio test (orange) and the Bayesian test (black) at different sample sizes ($x$-axis) at different ploidies (column facets) when either the null is true ("Null") or the alternative is true ("Alt"). The times for the Bayesian tests are the same for $K = 8$ and for large $n$ when $K = 6$ because we use simulation in those scenarios, and so any larger ploidy/sample size will result in the same runtime. The Bayesian test is slightly faster than the likelihood ratio test, but both are still fast enough to be computationally feasible for genome-wide applications.

**Procedure S3** Gibbs sampler and marginal likelihood estimation under the null of random mating using genotype likelihoods

**Input:** Genotype likelihoods $\ell_{ik}$ for $i = 1, 2, \ldots, n$ and $k = 0, 1, \ldots, K$, hyperparameter $\boldsymbol{\alpha} = (\alpha_0, \alpha_1, \ldots, \alpha_{K/2})$, $B$ sampling iterations, and $T$ burn-in iterations.
Initialize $\boldsymbol{p} \sim \text{Dirichlet}(\boldsymbol{\alpha})$
Initialize $\boldsymbol{q} \leftarrow \boldsymbol{p} * \boldsymbol{p}$
Initialize $\tilde{\boldsymbol{p}} \leftarrow \boldsymbol{p}$
Initialize $\tilde{\boldsymbol{q}} \leftarrow \tilde{\boldsymbol{p}} * \tilde{\boldsymbol{p}}$
Initialize $\tilde{L} \leftarrow \text{Dirichlet}(\boldsymbol{p}|\boldsymbol{\alpha}) \prod_{i=1}^{n} \sum_{k=0}^{K} \ell_{ik} q_k$
Initialize $\hat{\pi} \leftarrow 0$
**for** $j = 1, 2, \ldots, T + B$ **do**
    For $i = 1, 2, \ldots, n$, sample $z_i$ with probabilities $Pr(z_i = k) = \ell_{ik} q_k / \sum_{k=0}^{K} \ell_{ik} q_k$ for $k = 0, 1, \ldots, K$
    $x_k \leftarrow \sum_{i=1}^{n} 1(z_i = k)$ for $k = 0, 1, \ldots, K$
    Sample $[\boldsymbol{y}|\boldsymbol{x}, \boldsymbol{p}]$ using Procedure S2
    Sample $[\boldsymbol{p}|\boldsymbol{y}] \sim \text{Dirichlet}(\boldsymbol{y} + \boldsymbol{\alpha})$
    $\boldsymbol{q} \leftarrow \boldsymbol{p} * \boldsymbol{p}$
    **if** $(\text{Dirichlet}(\boldsymbol{p}|\boldsymbol{\alpha}) \prod_{i=1}^{n} \sum_{k=0}^{K} \ell_{ik} q_k > \tilde{L})$ **and** $(j \leq T)$ **then**
        $\tilde{\boldsymbol{p}} \leftarrow \boldsymbol{p}$
        $\tilde{\boldsymbol{q}} \leftarrow \tilde{\boldsymbol{p}} * \tilde{\boldsymbol{p}}$
        $\tilde{L} \leftarrow \text{Dirichlet}(\boldsymbol{p}|\boldsymbol{\alpha}) \prod_{i=1}^{n} \sum_{k=0}^{K} \ell_{ik} q_k$
    **else if** $j > T$ **then**
        $\hat{\pi} \leftarrow \hat{\pi} + \frac{1}{B} \text{Dirichlet}(\tilde{\boldsymbol{p}}|\boldsymbol{y} + \boldsymbol{\alpha})$
    **end if**
**end for**
Estimate marginal likelihood

$$\widehat{\Pr}(\boldsymbol{d}|H_0) = \frac{\text{Dirichlet}(\tilde{\boldsymbol{p}}|\boldsymbol{\alpha}) \prod_{i=1}^{n} \sum_{k=0}^{K} \ell_{ik} \tilde{q}_k}{\hat{\pi}(\tilde{\boldsymbol{p}}|\boldsymbol{d})} \tag{S80}$$

**Return:** Estimate of marginal likelihood $\widehat{\Pr}(\boldsymbol{d}|H_0)$.

**Procedure S4** Gibbs sampler and marginal likelihood estimation under the alternative of non-random mating using genotype likelihoods

**Input:** Genotype likelihoods $\ell_{ik}$ for $i = 1, 2, \ldots, n$ and $k = 0, 1, \ldots, K$, hyperparameter $\boldsymbol{\beta} = (\beta_0, \beta_1, \ldots, \beta_K)$, $B$ sampling iterations, and $T$ burn-in iterations.
Initialize $\boldsymbol{q} \sim \text{Dirichlet}(\boldsymbol{\beta})$
Initialize $\tilde{\boldsymbol{q}} \leftarrow \boldsymbol{q}$
Initialize $\tilde{L} \leftarrow \text{Dirichlet}(\boldsymbol{q}|\boldsymbol{\beta}) \prod_{i=1}^{n} \sum_{k=0}^{K} \ell_{ik} q_k$.
Initialize $\hat{\pi} \leftarrow 0$
**for** $j = 1, 2, \ldots, T + B$ **do**
  For $i = 1, 2, \ldots, n$, sample $z_i$ with probabilities $Pr(z_i = k) = \ell_{ik} q_k / \sum_{k=0}^{K} \ell_{ik} q_k$ for $k = 0, 1, \ldots, K$
  $x_k \leftarrow \sum_{i=1}^{n} 1(z_i = k)$ for $k = 0, 1, \ldots, K$
  Sample $[\boldsymbol{q}|\boldsymbol{x}] \sim \text{Dirichlet}(\boldsymbol{x} + \boldsymbol{\beta})$
  **if** $(\text{Dirichlet}(\boldsymbol{q}|\boldsymbol{\beta}) \prod_{i=1}^{n} \sum_{k=0}^{K} \ell_{ik} q_k > \tilde{L})$ **and** $(j \leq T)$ **then**
    $\tilde{\boldsymbol{q}} \leftarrow \boldsymbol{q}$
    $\tilde{L} \leftarrow \text{Dirichlet}(\boldsymbol{q}|\boldsymbol{\beta}) \prod_{i=1}^{n} \sum_{k=0}^{K} \ell_{ik} q_k$
  **else if** $j > T$ **then**
    $\hat{\pi} \leftarrow \hat{\pi} + \frac{1}{B} \text{Dirichlet}(\tilde{\boldsymbol{q}}|\boldsymbol{x} + \boldsymbol{\beta})$
  **end if**
**end for**
Estimate marginal likelihood

$$\widehat{\text{Pr}}(\boldsymbol{d}|H_0) = \frac{\text{Dirichlet}(\tilde{\boldsymbol{q}}|\boldsymbol{\beta}) \prod_{i=1}^{n} \sum_{k=0}^{K} \ell_{ik} \tilde{q}_k}{\hat{\pi}(\tilde{\boldsymbol{q}}|\boldsymbol{d})} \tag{S81}$$

**Return:** Estimate of marginal likelihood $\widehat{\text{Pr}}(\boldsymbol{d}|H_0)$.

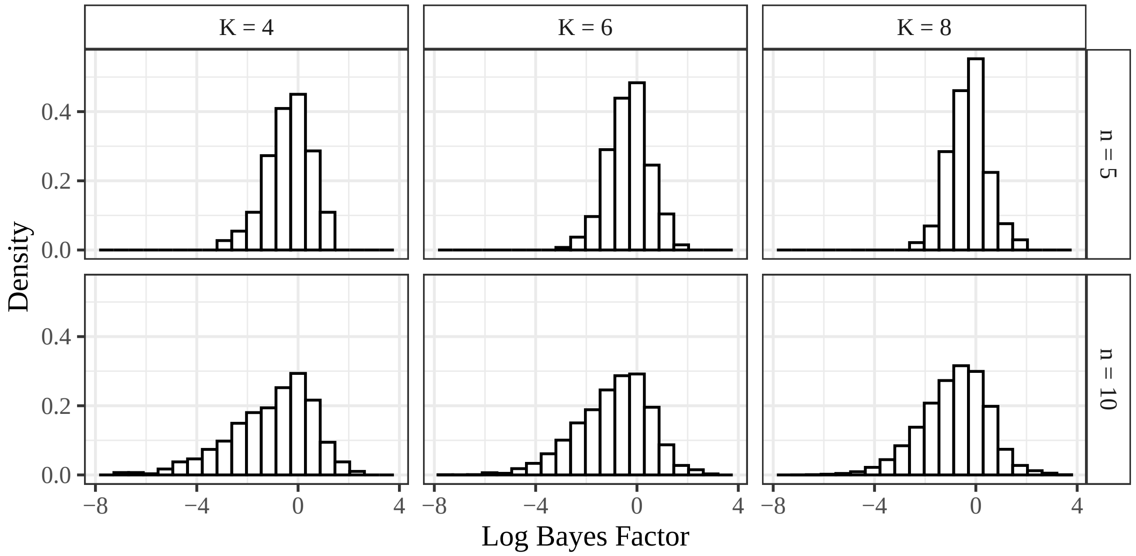


Figure S2: Histogram of all log Bayes factors for $n = 5$ and 10 (row facets) and ploidies $K = 4$, 6, and 8 (column facets).

| Ploidy | $n$ | log BF | $x_0$ | $x_1$ | $x_2$ | $x_3$ | $x_4$ | $x_5$ | $x_6$ | $x_7$ | $x_8$ |
|---|---|---|---|---|---|---|---|---|---|---|---|
| 4 | 5 | 1.20 | 0 | 0 | 0 | 1 | 4 | - | - | - | - |
| 4 | 10 | 2.16 | 0 | 0 | 0 | 2 | 8 | - | - | - | - |
| 6 | 5 | 1.54 | 0 | 0 | 0 | 0 | 0 | 1 | 4 | - | - |
| 6 | 10 | 2.87 | 0 | 0 | 0 | 0 | 0 | 2 | 8 | - | - |
| 8 | 5 | 1.74 | 3 | 2 | 0 | 0 | 0 | 0 | 0 | 0 | 0 |
| 8 | 10 | 3.37 | 6 | 3 | 1 | 0 | 0 | 0 | 0 | 0 | 0 |
| 4 | 5 | -2.72 | 2 | 0 | 0 | 3 | 0 | - | - | - | - |
| 4 | 10 | -6.82 | 3 | 0 | 0 | 7 | 0 | - | - | - | - |
| 6 | 5 | -2.74 | 0 | 3 | 0 | 0 | 0 | 2 | 0 | - | - |
| 6 | 10 | -7.69 | 0 | 5 | 0 | 0 | 0 | 5 | 0 | - | - |
| 8 | 5 | -2.50 | 0 | 2 | 0 | 0 | 0 | 0 | 0 | 3 | 0 |
| 8 | 10 | -7.18 | 0 | 5 | 0 | 0 | 0 | 0 | 0 | 5 | 0 |

Table S2: Small sample datasets with highest and lowest Bayes factors by ploidy and sample size. Very large Bayes factors tend to be unimodal, whereas very small Bayes factors tend to have a couple prominent modes.

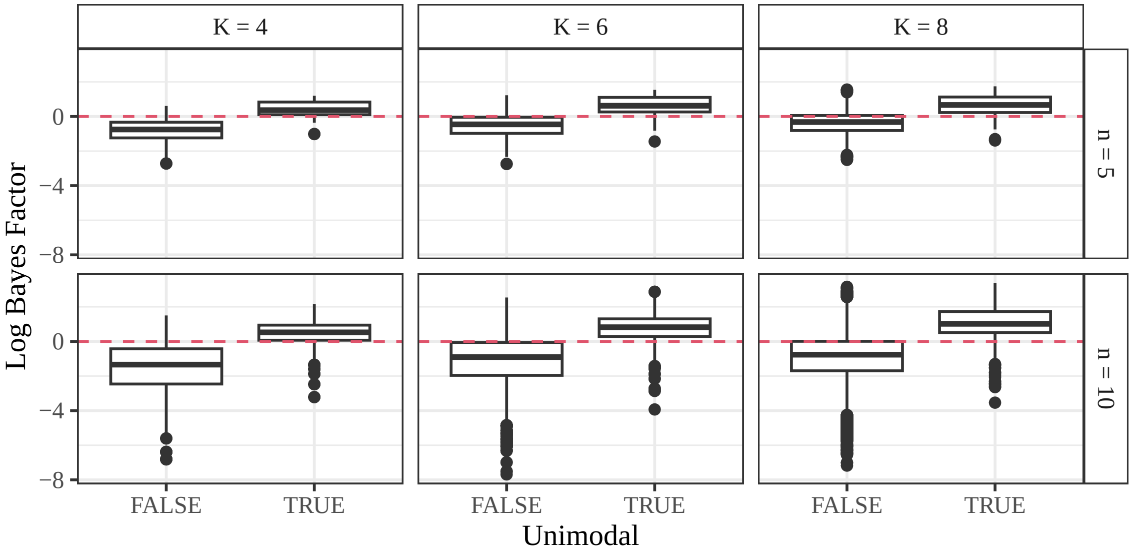


Figure S3: Boxplots of log Bayes factors ($y$-axis) by unimodality status of $\boldsymbol{x}$ ($x$-axis), stratified by ploidy (column facets) and sample size (row facets). Unimodal $\boldsymbol{x}$ tends to result in positive log Bayes factors, while non-unimodal $\boldsymbol{x}$ tends to result in negative log Bayes factors.

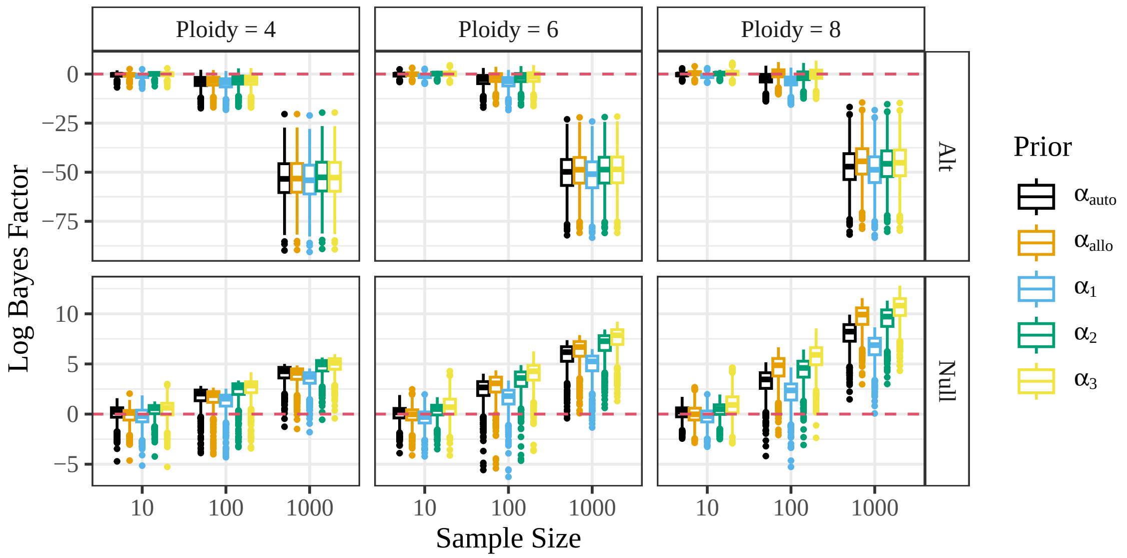


Figure S4: Log Bayes factors ($y$-axis) at different sample sizes ($x$-axis) and different prior distributions (colors) at different ploidies (column facets) when either the null is true ("Null") or the alternative is true ("Alt"). The Bayes factor is consistent for any prior selection. Even moderate deviations from the default prior ($\boldsymbol{\alpha}_0$) result in only modest differences in the Bayes factors.

| SNP | $x_0$ | $x_1$ | $x_2$ | $x_3$ | $x_4$ | log BF |
|---|---|---|---|---|---|---|
| Atr_20529-52 | 0 | 6 | 2 | 1 | 10 | -5.9 |
| Atr_33123-54 | 1 | 5 | 4 | 0 | 9 | -4.7 |
| Atr_26310-63 | 0 | 6 | 4 | 1 | 8 | -4.2 |
| Atr_7817-58 | 2 | 5 | 2 | 2 | 8 | -3.1 |
| Atr_13476-65 | 0 | 4 | 2 | 2 | 11 | -2.6 |
| Atr_13917-71 | 1 | 16 | 2 | 0 | 0 | -2.6 |
| Atr_27367-69 | 1 | 6 | 1 | 6 | 5 | -2.4 |
| Atr_10998-72 | 1 | 6 | 1 | 8 | 3 | -2.3 |
| Atr_37739-28 | 4 | 2 | 2 | 5 | 6 | -2.1 |
| Atr_48754-60 | 3 | 1 | 2 | 7 | 6 | -2.0 |

Table S3: SNPs from the white sturgeon data of Section 3.3 that have log Bayes factors below -2. Each $x_k$ is the number of individuals with genotype $k$ at that SNP. The SNPs with the lowest three Bayes factors appear to be anomalous.

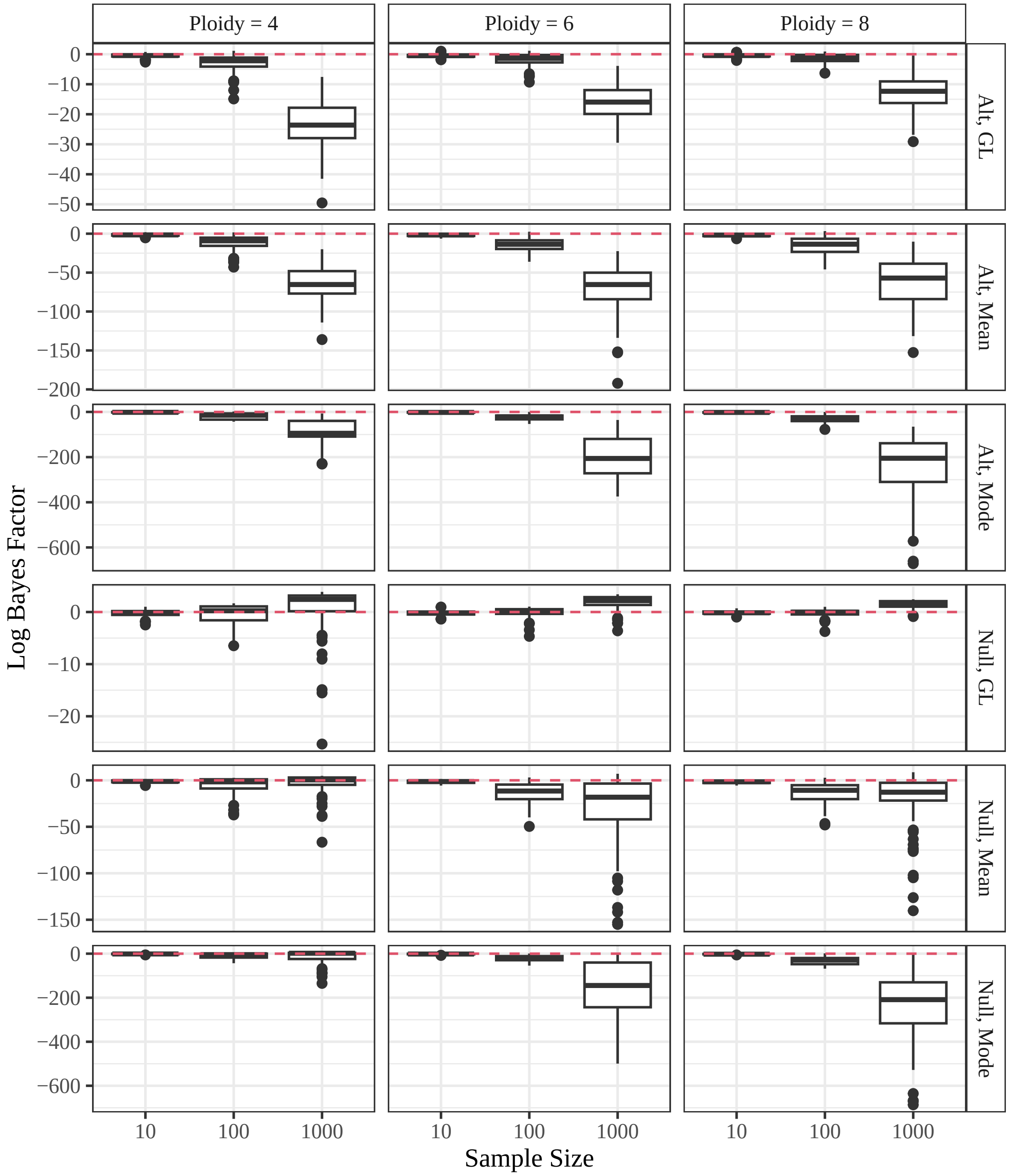


Figure S5: Log Bayes factors ($y$-axis) by sample size ($x$-axis) when either the null is satisfied ("Null") or the alternative is satisfied ("Alt"), and when using either the genotype likelihood approach from Section 2.2 ("GL") or when using the known genotype approach from Section 2.1 and an estimate of $\boldsymbol{x}$ ("Mean" or "Mode"). The two approaches to estimate $\boldsymbol{x}$ were either to sum the posterior probabilities ("Mean") or to tabulate the posterior modes ("Mode"). Bayes factors should increase when the null is satisfied and should decrease when the alternative is satisfied, but only do so when using the genotype likelihood approach.

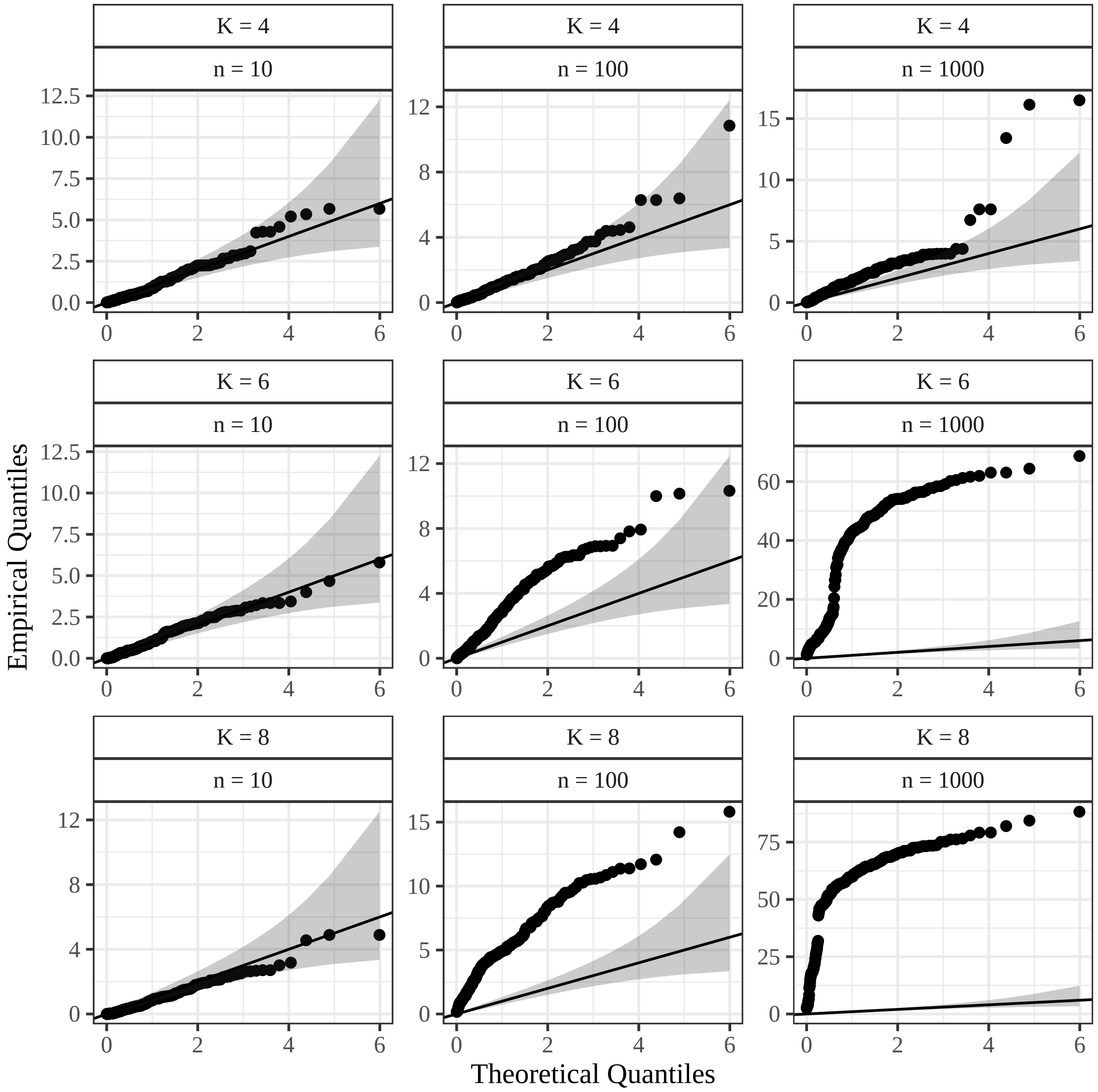


Figure S6: QQ-plots, when the null is true, of $p$-values of the likelihood ratio test for random mating against the uniform distribution, on the $-\log$ scale, with the 95% simultaneous confidence bands from Aldor-Noiman et al. [2013], stratified by ploidy (row facets) and sample size (column facets). These $p$-values come from the simulations of Section 3.2 where genotypes are not known, and the likelihood parameters are estimated during genotyping with the proportional normal prior class. The likelihood ratio test assumes known genotypes, and so genotypes were estimated using the posterior modes. Since the null is satisfied for these simulations, the $p$-values should follow a uniform distribution. However, because the likelihood ratio test assumes known genotypes, there are severe deviations from the uniform distribution that worsen with larger sample sizes. This would result in an excess of false positives.

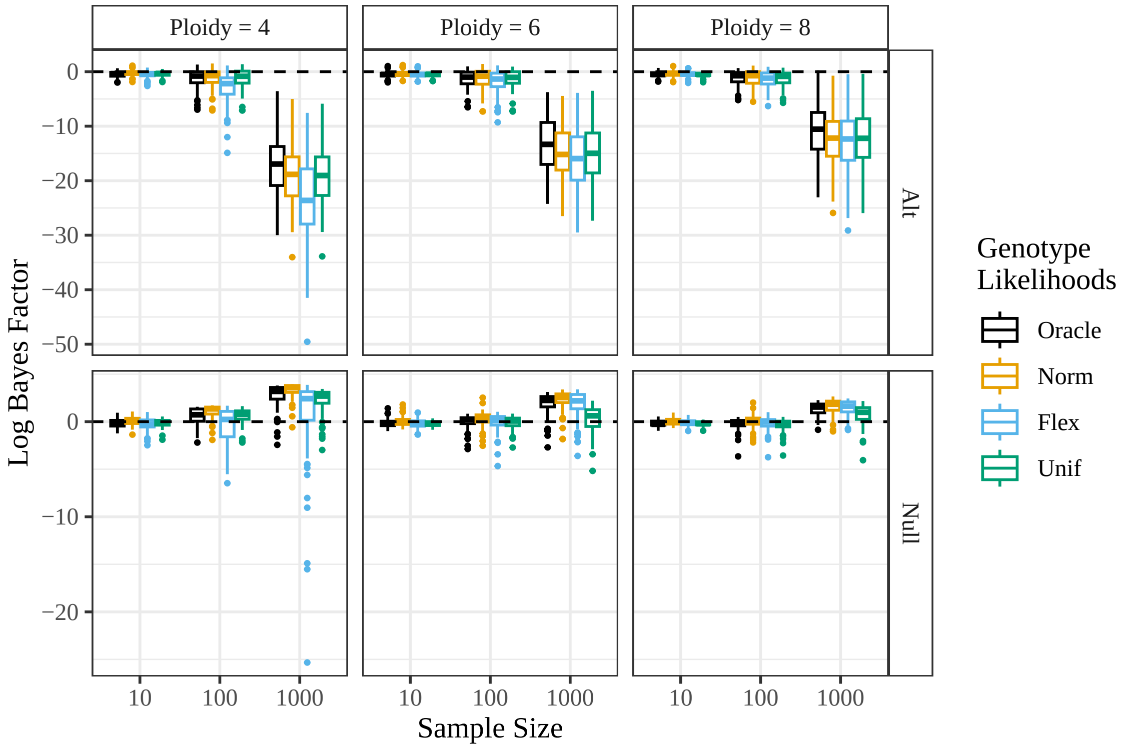


Figure S7: Log Bayes factors ($y$-axis) by sample size ($x$-axis) when either the null is satisfied ("Null") or the alternative is satisfied ("Alt"), when using the genotype likelihood approach from Section 2.2. Genotype likelihoods were generated under different parameterizations of `updog` [Gerard et al., 2018]: using the true likelihood parameters ("Oracle"), using the proportional normal prior class [Gerard and Ferrão, 2019] ("Norm"), using the general categorical prior class ("Flex"), or using the discrete uniform distribution ("Unif"). Bayes factors should increase when the null is satisfied and should decrease when the alternative is satisfied. All approaches to genotyping accomplish this, though the uniform prior class performs the worst. This indicates that our methods are very robust to genotype likelihood miscalibration.

| | | | | Observed/Expected | | | | | | |
|---|---|---|---|---|---|---|---|---|---|---|
| SNP | Pr(Mis) | Log BF | Parent | 0 | 1 | 2 | 3 | 4 | 5 | 6 |
| Itr_sc035768.1:536_G/C | 0.07 | -14.17 | 5 | 9/0 | 1/0 | 0/0 | 0/0 | 27/35 | 63/70 | 40/35 |
| Itr_sc002687.1:16048_G/A | 0.08 | -10.88 | 5 | 0/0 | 0/0 | 0/0 | 0/0 | 14/35 | 98/70 | 28/35 |
| Itr_sc000111.1:443840_A/C | 0.07 | -9.09 | 5 | 6/0 | 1/0 | 1/0 | 0/0 | 34/35 | 53/70 | 46/35 |
| Itr_sc003300.1:9036_A/G | 0.41 | -8.69 | 3 | 30/0 | 5/6 | 29/35 | 41/58 | 27/35 | 8/6 | 1/0 |
| Itr_sc002687.1:16006_A/G | 0.13 | -8.20 | 1 | 37/35 | 71/70 | 29/35 | 0/0 | 0/0 | 1/0 | 3/0 |

Table S4: Five SNPs from the sweet potato data of Section 3.4 with log Bayes factors less than -5. The estimated proportion of individuals mis-genotyped and the estimated parental genotype were provided by `updog` [Gerard et al., 2018, Gerard and Ferrão, 2019], and the log Bayes factor was calculated using the method of Section 2.2. The right half of the table contains the observed/expected counts for the genotypes 0 through 6. The expected counts were calculated assuming the parental genotype is correct using a convolution of hypergeometric distributions Serang et al. [2012].

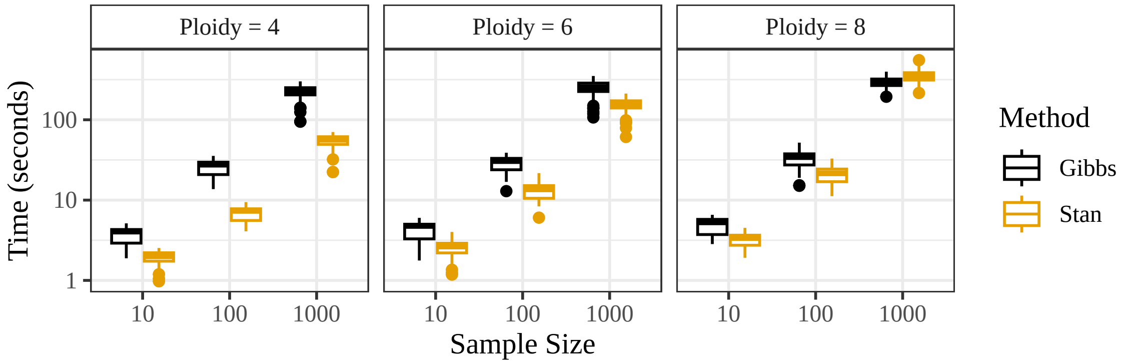


Figure S8: Time of evaluation ($y$-axis) for Stan (Section 2.2) or the Gibbs samplers from Appendix S3 for different sample sizes ($x$-axis) and different ploidies (facets). The number of samples for Stan and the Gibbs sampler were chosen so that they take about the same amount of time when $n = 1000$ and $K = 8$.

| | S1 Test | |
|---|---|---|
| RM Test | log(BF) $> 0$ | log(BF) $< 0$ |
| log(BF) $> 0$ | 4931 | 123 |
| log(BF) $< 0$ | 3 | 17 |

Table S5: Contingency table for whether the random mating test (rows) or S1 test (columns) provide evidence for the null (log(BF) $> 0$) or the alternative (log(BF) $< 0$).

| SNP | $\hat{x}_0$ | $\hat{x}_1$ | $\hat{x}_2$ | $\hat{x}_3$ | $\hat{x}_4$ | $\hat{x}_5$ | $\hat{x}_6$ | PG | RM | S1 | S1 New |
|---|---|---|---|---|---|---|---|---|---|---|---|
| Itr_sc000033.1:181645_C/A | 1 | 0 | 2 | 0 | 29 | 72 | 38 | 5 | 3.8 | -27.3 | 13.3 |
| Itr_sc000035.1:22200_C/T | 1 | 0 | 1 | 0 | 38 | 71 | 30 | 5 | 3.8 | -24.8 | 11.2 |
| Itr_sc000133.1:223799_G/A | 2 | 1 | 0 | 0 | 30 | 68 | 40 | 5 | 1.3 | -47.2 | 6.6 |
| Itr_sc000298.1:15705_A/T | 1 | 0 | 0 | 0 | 32 | 77 | 32 | 5 | 2.6 | -21.4 | 12.9 |
| Itr_sc001683.1:13298_G/A | 1 | 0 | 0 | 0 | 31 | 78 | 31 | 5 | 2.6 | -21.9 | 11.5 |
| Itr_sc001683.1:13421_T/A | 1 | 0 | 0 | 0 | 31 | 78 | 31 | 5 | 2.4 | -22.7 | 12.0 |
| Itr_sc004380.1:5581_C/G | 2 | 0 | 0 | 0 | 33 | 72 | 35 | 5 | 0.3 | -38.8 | 13.2 |
| Itr_sc011872.1:11711_C/T | 2 | 0 | 0 | 0 | 29 | 70 | 41 | 5 | 0.7 | -26.6 | 8.4 |

Table S6: Counts of maximum *a posteriori* genotype estimates ($x_0, \ldots, x_6$) and estimated parental genotype ("PG") for a random sample of 8 SNPs where the test for random mating (Section 2.2) produces positive log Bayes factors ("RM"), and the test for S1 proportions (Section S7) produces negative log Bayes factors ("S1"). Parental genotype estimates are all 5, which indicates that genotypes 0 through 3 should be impossible, but we do have some individuals with those genotype estimates. When we remove those individuals, the S1 test now produces positive log Bayes factors ("S1 New"). This indicates the S1 test is very sensitive to violations in its assumptions.

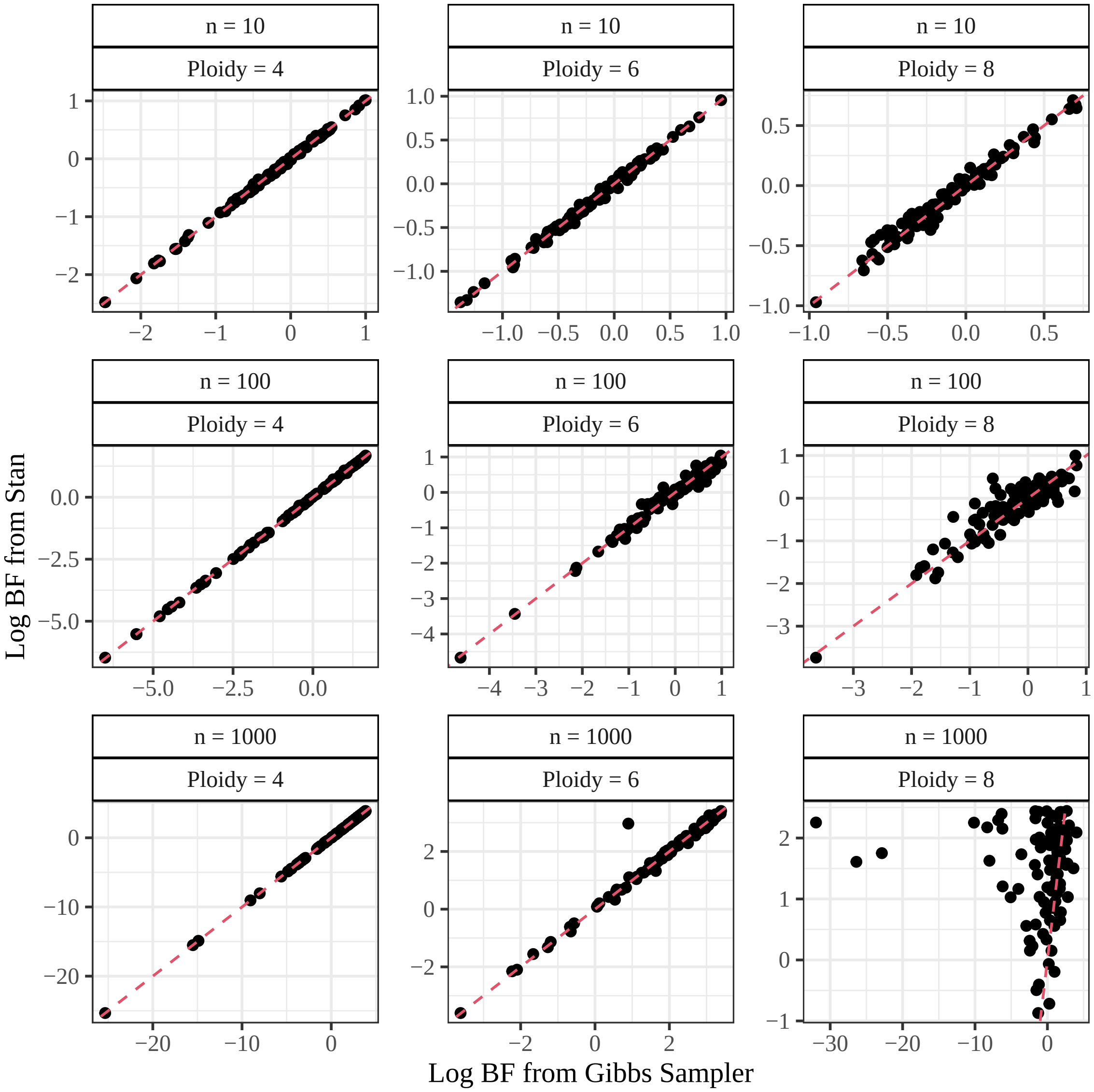


Figure S9: Bayes factor calculations using Stan ($y$-axis) (Section 2.2) versus the Gibbs samplers from Appendix S3 ($x$-axis) when the null is satisfied, for different sample sizes and different ploidies. Estimates line up except for large samples of octoploids, where the Gibbs sampler had poor mixing.

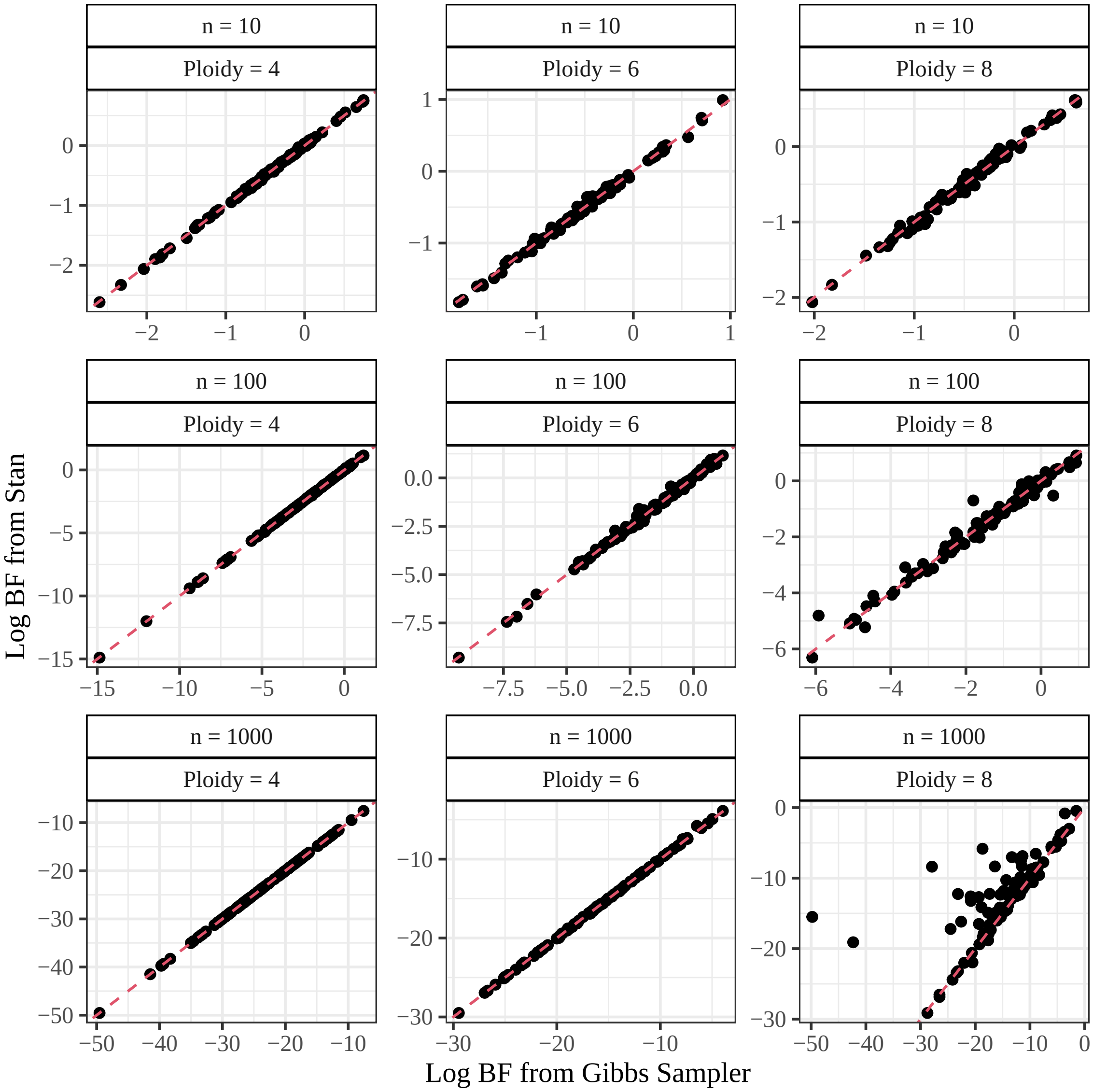


Figure S10: Bayes factor calculations using Stan ($y$-axis) (Section 2.2) versus the Gibbs samplers from Appendix S3 ($x$-axis) when the alternative is satisfied, for different sample sizes and different ploidies. Estimates line up except for large samples of octoploids, where the Gibbs sampler had poor mixing.

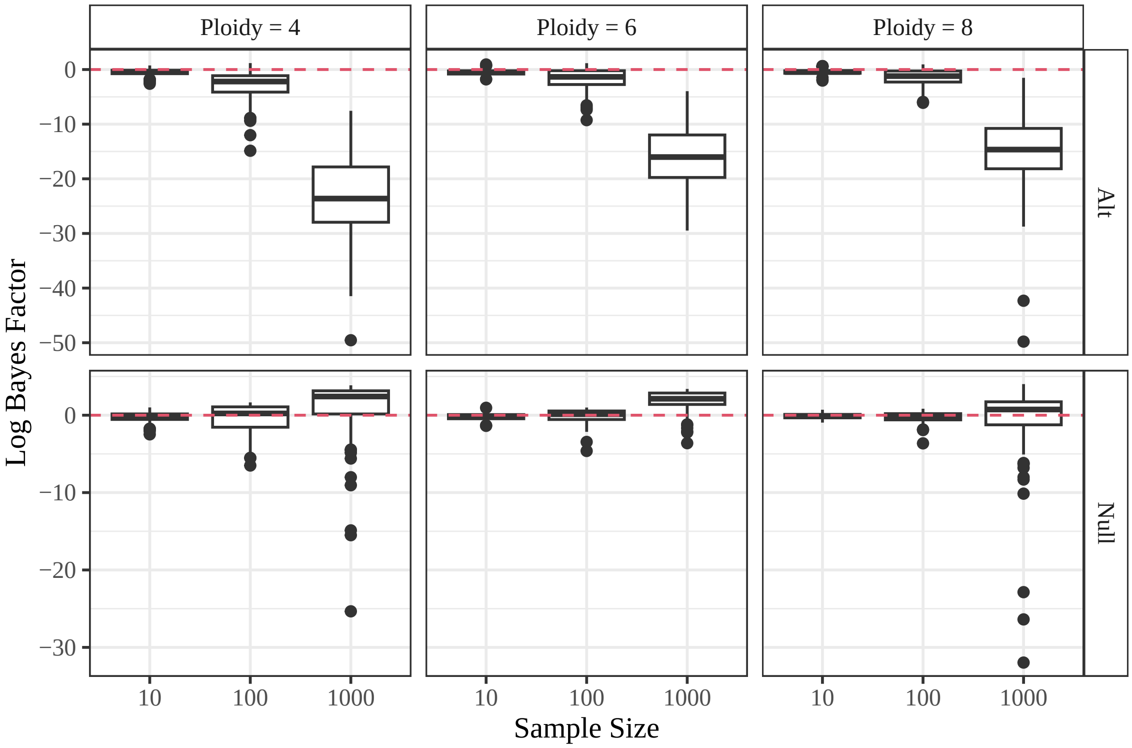


Figure S11: Log Bayes factor ($y$-axis) calculated using the Gibbs samplers from Appendix S3 when either the null is fulfilled ("Null") or the alternative is fulfilled ("Alt") at different sample sizes ($x$-axis) and different ploidies (column facets). Bayes factors should decrease under the alternative and increase under the null, and they do for ploidies of $K = 4$ and 6. However, for $K = 8$, the poor mixing the Gibbs sampler resulted in inaccurate calculations of the Bayes factor and the Bayes factor does not increase under the null. This is why we resorted to using Stan in Section 2.2.

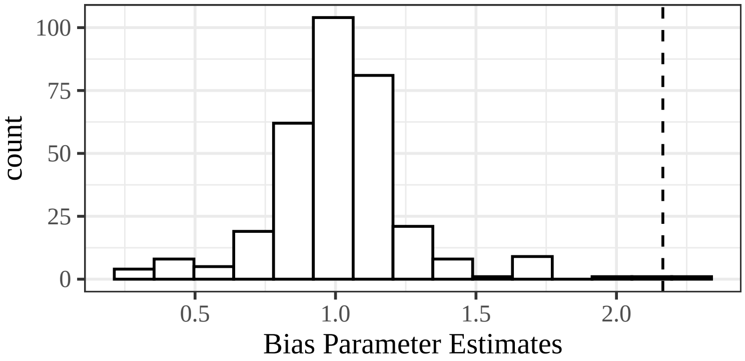


Figure S12: Histogram of bias parameter estimates from an `updog` fit [Gerard et al., 2018, Gerard and Ferrão, 2019] on the white sturgeon data from Section 3.3. Values above 1 indicate bias towards the alternative allele, and values below 1 indicate bias towards the reference allele. The vertical dashed line is the bias estimate at the local mode fit of SNP Atr_20529-52.

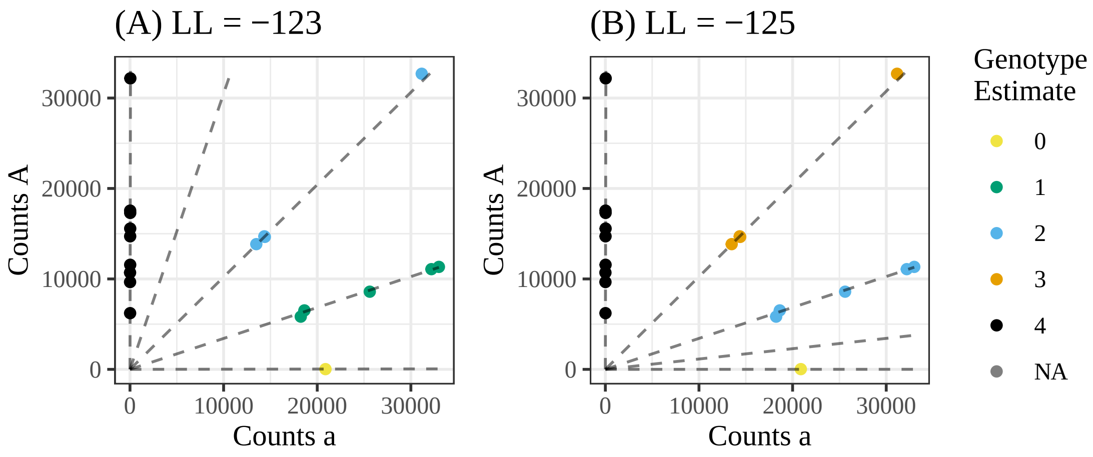


Figure S13: Genotype plots [Gerard et al., 2018] for two fits on SNP Atr_33123-54 from the white sturgeon data of Section 3.3. See Figure 5 for a description of genotype plots. The left plot **(A)** is the global mode with a log-likelihood of -123, and the right plot **(B)** is a local mode with a log-likelihood of -125. The test for random mating using the global mode of $\boldsymbol{x} = (1, 5, 4, 0, 9)$ yields a log Bayes factor of -4.7, while the test for random mating using the local mode of $\boldsymbol{x} = (1, 0, 5, 4, 9)$ yields a log Bayes factor of 0.8. The global mode is thus likely a genotyping error.

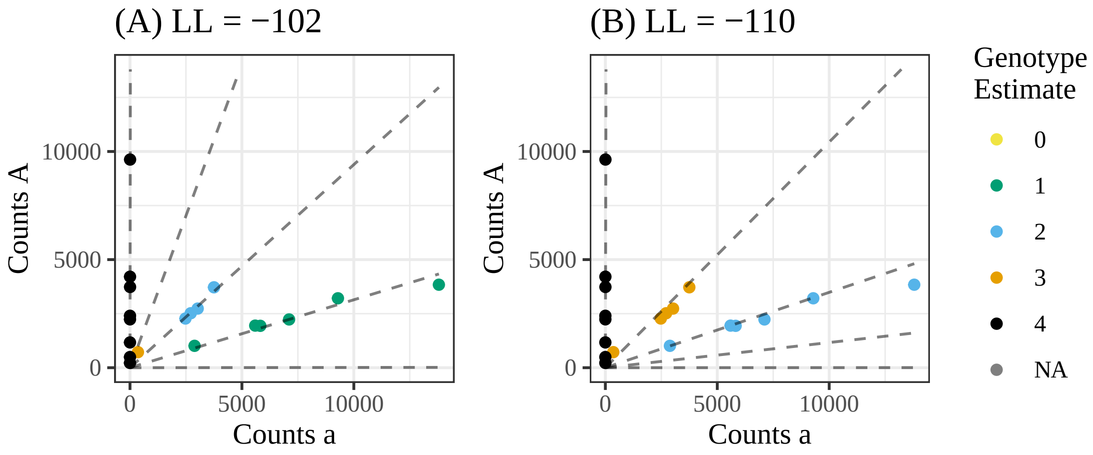


Figure S14: Genotype plots [Gerard et al., 2018] for two fits on SNP Atr_26310-63 from the white sturgeon data of Section 3.3. See Figure 5 for a description of genotype plots. The left plot **(A)** is the global mode with a log-likelihood of -102, and the right plot **(B)** is a local mode with a log-likelihood of -110. The test for random mating using the global mode of $\boldsymbol{x} = (0, 6, 4, 1, 8)$ yields a log Bayes factor of -4.2, while the test for random mating using the local mode of $\boldsymbol{x} = (0, 0, 6, 5, 8)$ yields a log Bayes factor of 1.2. The global mode is thus likely a genotyping error.

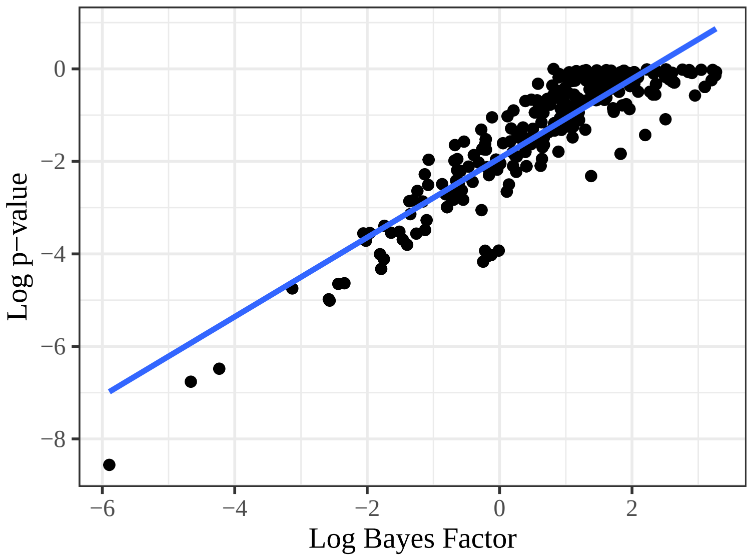


Figure S15: Scatterplot of log Bayes factors ($x$-axis) and log $p$-values ($y$-axis) from the white sturgeon data from Section 3.3, with their ordinary least squares line.

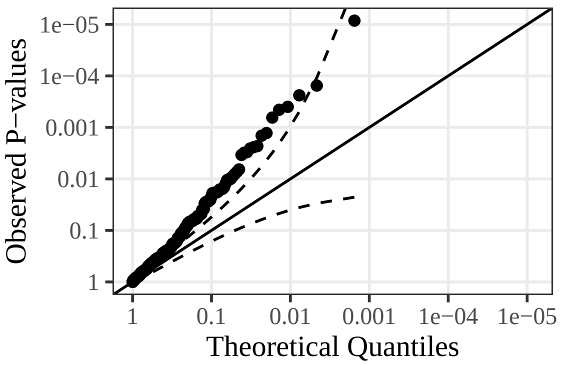


Figure S16: Quantile-quantile plots, on the $-\log_{10}$-scale, of the $p$-values from the $U$-statistic test for equilibrium from Gerard [2022] applied to the white sturgeon data of Section 3.3. 95% confidence bands, calculated using the procedure from Aldor-Noiman et al. [2013], are provided. The $p$-values are highly non-uniformly distributed.

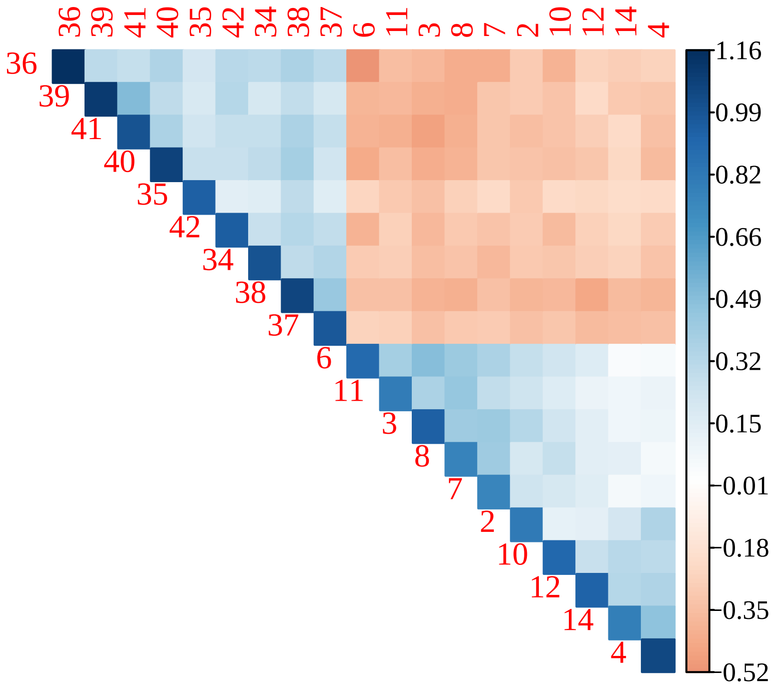


Figure S17: Pairwise relatedness measures of Ashraf et al. [2016] (color), calculated on all 19 samples (rows and columns) of the white sturgeon data of Section 3.3. Samples were ordered by the angular ordering of the eigenvectors [Friendly, 2002] to highlight closely related groups. The plot was made using `corrplot` [Wei and Simko, 2021].

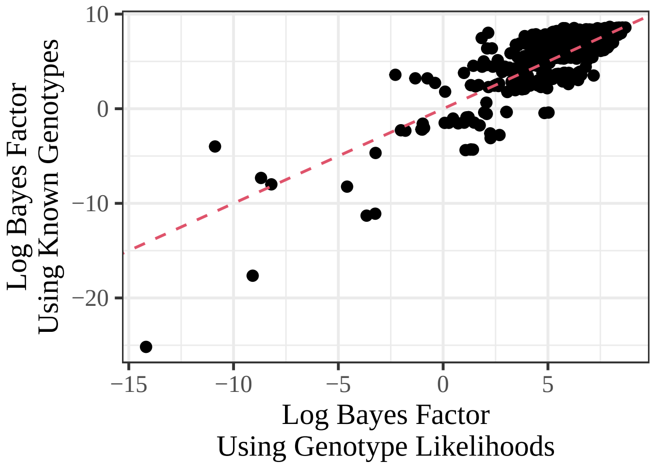


Figure S18: Scatterplot of log Bayes factors using either genotype likelihoods ($x$-axis) or assuming the sum of the posterior probabilities are the known genotypes ($y$-axis) for the sweet potato data from Section 3.4. The low correlation indicates that we should use the genotype likelihood approach for these data.

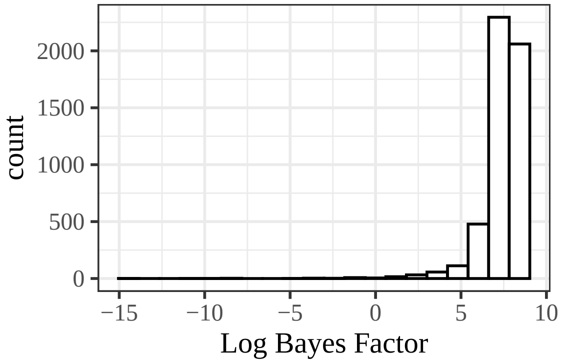


Figure S19: Histogram of the log Bayes factors using the test from Section 2.2 on the sweet potato data from Section 3.4.

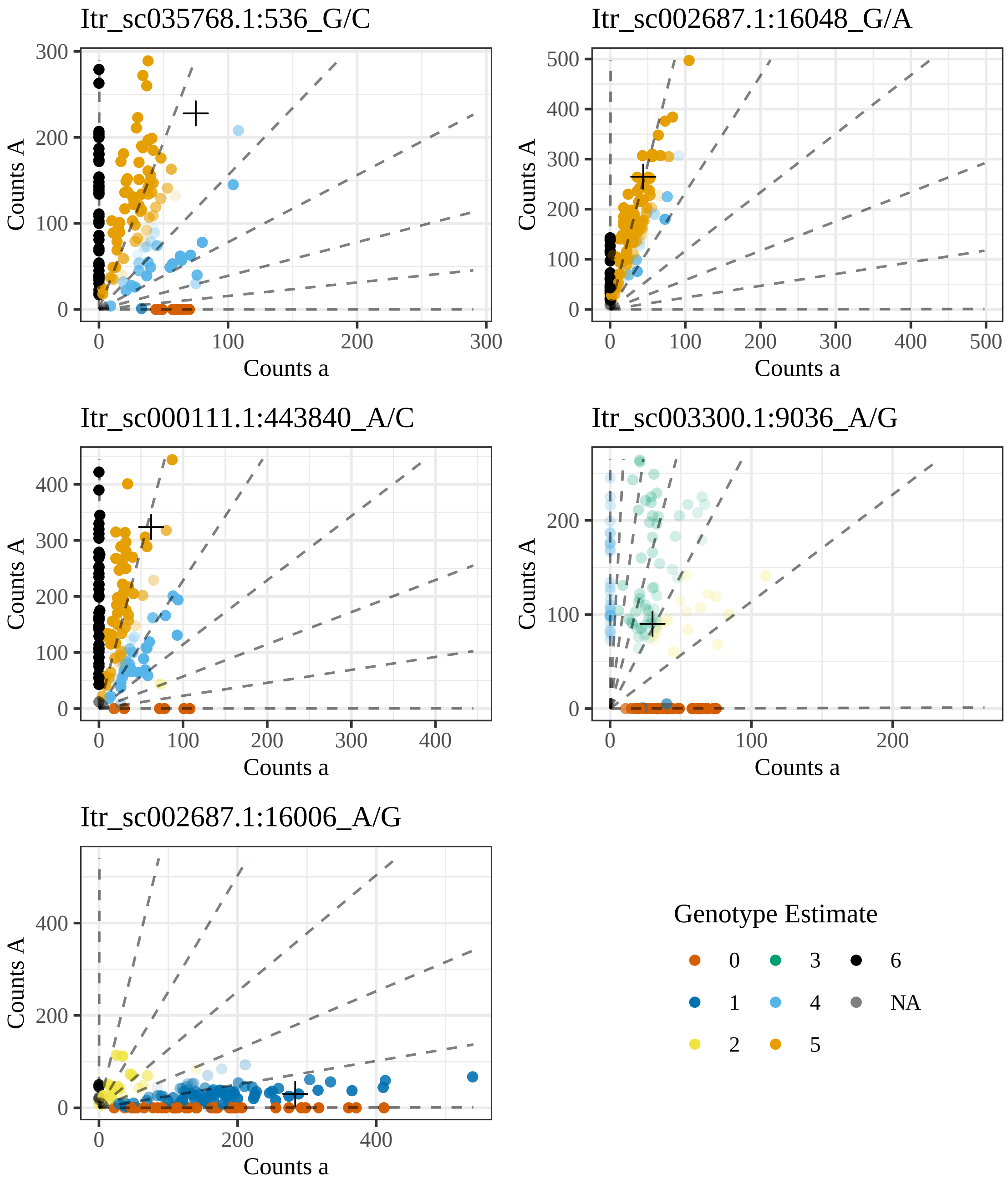


Figure S20: Genotype plots for the 5 loci that have log Bayes factors less than -5, for the test of random mating, from the sweet potato data of Section 3.4. See Figure 5 for a description of genotype plots.

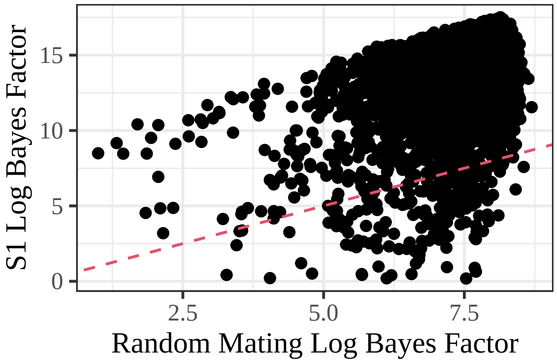


Figure S21: Scatterplot of the log Bayes factors using either the null of random mating ($x$-axis) or the null of S1 segregation proportions ($y$-axis), applied to the sweet potato data from section 3.4 where both null hypotheses are satisfied, for loci where both log Bayes factors are greater than zero. The dashed line is the $y = x$ line. The S1 test tends to produce larger Bayes factors, being more sure of the null hypothesis.

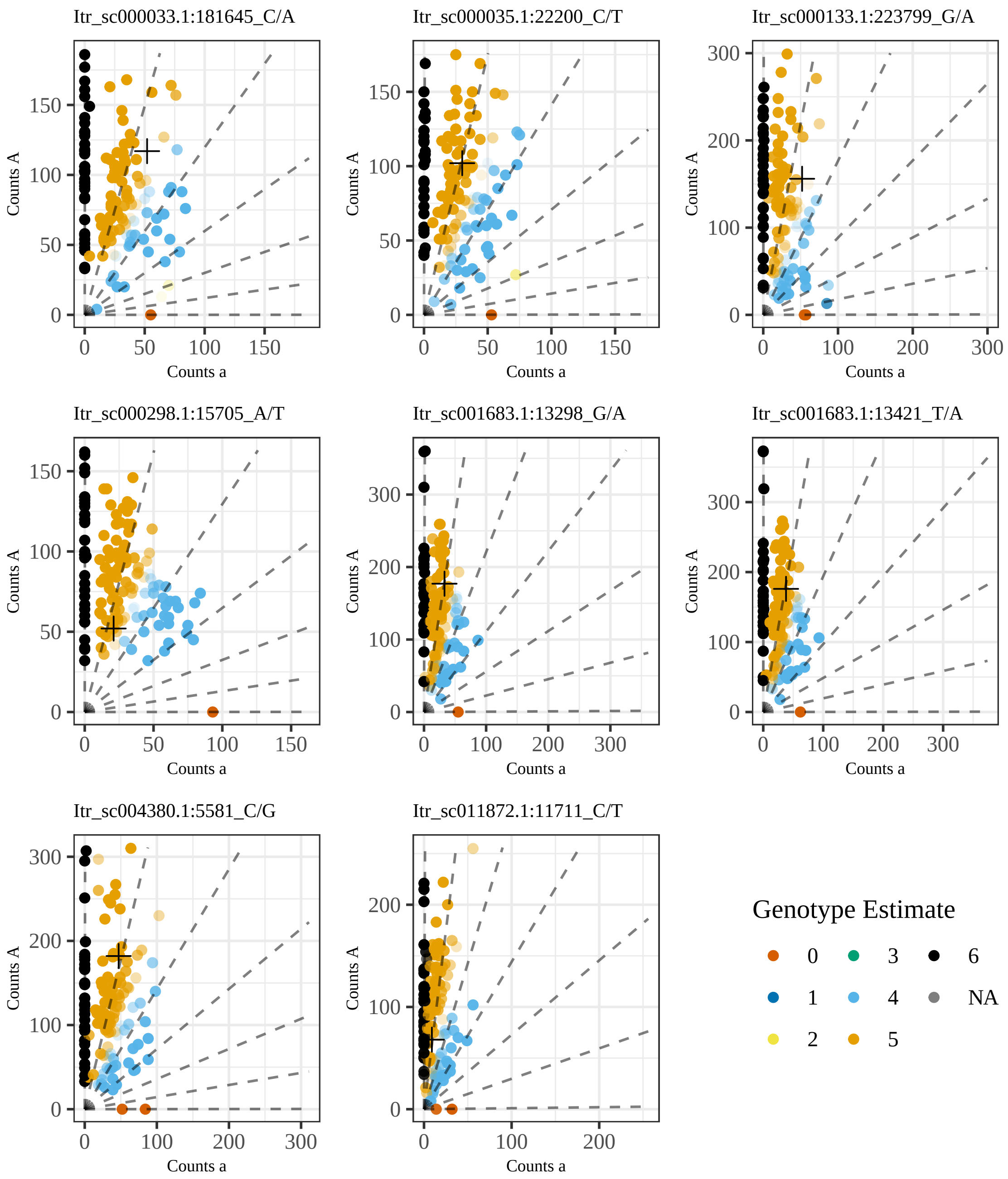


Figure S22: Genotype plots for a random sample of eight SNPs from the sweet potato data of Section 3.4 where the test for random mating (Section 2.2) produces positive log Bayes factors, and the test for S1 proportions (Section S7) produces negative log Bayes factors. Each SNP is relatively well behaved except for two or three individuals. See Figure 5 for a description of genotype plots.

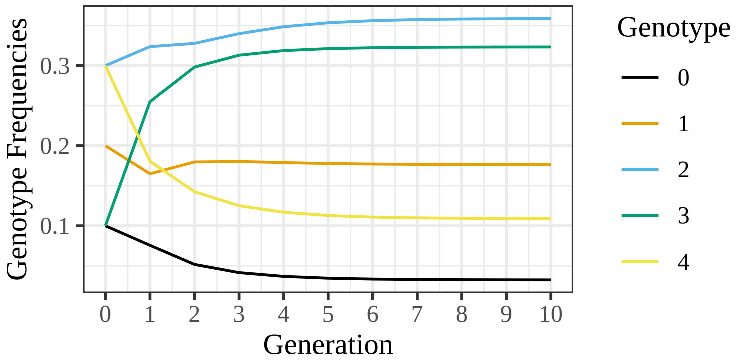


Figure S23: Frequencies ($y$-axis) of allopolyploid genotypes (color) over multiple generations of random mating ($x$-axis) when the joint distribution of subgenome genotypes starts at (S73). HWP are only approached asymptotically.
